# Coordinated and distinct LD-transpeptidase-independent roles of peptidoglycan carboxypeptidases DacC and DacA in stress adaptation and cell shape maintenance

**DOI:** 10.1101/2022.09.06.506770

**Authors:** Umji Choi, Si Hyoung Park, Han Byeol Lee, Chang-Ro Lee

**Affiliations:** Department of Biological Sciences, Myongji University, Yongin, Gyeonggido 449-728, Republic of Korea; The Natural Science Research Institute, Myongji University, Yongin, Gyeonggido 449-728, Republic of Korea

## Abstract

Peptidoglycan (PG) is an essential bacterial architecture pivotal for shape maintenance and adaptation to osmotic stress. Although PG synthesis and modification are tightly regulated under harsh environmental stresses, few related mechanisms have been investigated. In this study, we aimed to investigate the coordinated and distinct roles of the PG carboxypeptidases DacC and DacA, in adaptation to alkaline and salt stresses and shape maintenance in *Escherichia coli*. We found that DacC is an alkaline PG carboxypeptidase, whose enzyme activity and protein stability are significantly enhanced under alkaline stress. Both DacC and DacA were required for bacterial growth under alkaline stress, whereas only DacA was required for the adaptation to salt stress. Under normal growth conditions, only DacA was necessary for cell shape maintenance, while under alkaline stress conditions, both DacA and DacC were necessary for cell shape maintenance, but their roles were distinct. Notably, all these roles of DacC and DacA were independent of LD-transpeptidases, which are necessary for the formation of PG 3-3 crosslinks and covalent bonds between PG and the outer membrane lipoprotein Lpp. Instead, DacC and DacA interacted with penicillin-binding proteins (PBPs), DD-transpeptidases, mostly in a C-terminal domain-dependent manner, and these interactions were necessary for most of their roles. Collectively, our results demonstrate the coordinated and distinct novel roles of PG carboxypeptidases in stress adaptation and shape maintenance and provide novel insights into the cellular functions of PG carboxypeptidases associated with PBPs.

## Introduction

Peptidoglycan (PG) is a pivotal bacterium-specific architecture found in most Gram-negative and Gram-positive bacteria. PG not only determines bacterial shape but also plays an important role in providing protection aginst environmental stresses, such as turgor pressure (Vollmer and Bertsche, 2008; Rohs and Bernhardt, 2021). PG is a mesh-like polymer composed of long linear polysaccharide chains formed by alternating connections between two amino sugar derivatives, *N*-acetylglucosamine and *N*-acetylmuramic acid (Vollmer and Bertsche, 2008). The crosslinks between polysaccharide chains are formed by covalent bonds between the peptide chains, which are attached to *N*-acetylmuramic acid (Vollmer and Bertsche, 2008). The linear polysaccharide chain is formed by periplasmic enzymes with glycosyltransferase activity, such as the shape, elongation, division, and sporulation (SEDS) family of proteins and class A penicillin-binding proteins (PBPs) (Dorr, 2021; Rohs and Bernhardt, 2021). The crosslinks between polysaccharide chains are facilitated by periplasmic enzymes with DD-transpeptidase activity, such as class A or B PBP (aPBP or bPBP, respectively), which results in the formation of 4-3 crosslinks (Rohs and Bernhardt, 2021). The crosslinks between polysaccharide chains can also be formed by LD-transpeptidases, which results in the formation of 3-3 crosslinks (Hugonnet *et al.*, 2016). In most bacteria, the vast majority (>90%) of PG crosslinks are the 4-3 crosslinks, but in a few bacteria, such as *Mycobacterium tuberculosis*, the portion of 3-3 crosslinks is considerably high, accounting for >60% of the PG crosslinks (Baranowski *et al.*, 2018).

Crosslinked PG in the periplasm is degraded by diverse PG hydrolases, namely, amidases, lytic transglycosylases, endopeptidases, and carboxypeptidases (Vollmer *et al.*, 2008; van Heijenoort, 2011; Vermassen *et al.*, 2019). PG amidases that hydrolyze the lactylamide bond between *N*-acetylmuramic acid and the peptide chain are associated with cytokinetic ring contraction at the division site (Uehara *et al.*, 2010; Mueller *et al.*, 2021). Lytic transglycosylases that cleave the β-1,4-glycosidic bond between *N*-acetylglucosamine and *N*-acetylmuramic acid are known to play important roles in PG quality control (Cho *et al.*, 2014), termination of PG glycan polymerization (Yunck *et al.*, 2016; Bohrhunter *et al.*, 2021; Sassine *et al.*, 2021), and establishment of flagella and type VI secretion architectures (Nambu *et al.*, 1999; Santin and Cascales, 2017). PG endopeptidases that cleave within the cross-bridged peptide chains are known to function as space makers that cleave the crosslinked peptide chains for the insertion of newly synthesized PG strands (Singh *et al.*, 2012; Lai *et al.*, 2017; Chodisetti and Reddy, 2019; Park *et al*., 2020). All PG hydrolases showed strong redundancy. For example, *Escherichia coli* has nine enzymes that exhibit lytic transglycosylase activity in the periplasm (Bohrhunter *et al.*, 2021; Sassine *et al.*, 2021) and eight enzymes with PG endopeptidase activity (Chodisetti and Reddy, 2019; Park *et al.*, 2020). Although the physiological significance of PG hydrolase redundancy has not yet been fully elucidated, it is thought to enable the maintenance of PG synthesis and degradation under harsh extracellular stresses (Peters *et al.*, 2016; Mueller *et al.*, 2019; Park *et al*., 2020; Mueller *et al.*, 2021). Like MepK, which exclusively cleaves the 3-3 crosslinks, an enzyme with a narrow substrate specificity has also been reported (Chodisetti and Reddy, 2019).

PG carboxypeptidases remove the C-terminal fifth D-alanine (D-Ala) of the peptide chains (Vermassen *et al.*, 2019). After the removal of the fifth D-Ala, the remaining four amino acids of the peptide chain can be crosslinked with the adjacent peptide chain by LD-transpeptidases (LdtD and LdtE), or covalently attached to an abundant outer membrane (OM) lipoprotein Lpp (Braun’s lipoprotein) by other LD-transpeptidases (LdtA, LdtB, and LdtC) (Hugonnet *et al.*, 2016; Vermassen *et al.*, 2019). Although the roles of PG carboxypeptidases associated with LD-transpeptidases have been extensively studied, their roles in cell shape maintenance and PG synthesis regulation are poorly understood (Dorr, 2021). Seven proteins have PG carboxypeptidase activity in *E. coli* (van Heijenoort, 2011; Vermassen *et al.*, 2019). DacA (also called PBP5) is the most abundant PG carboxypeptidase and is expressed primarily during early exponential growth in *E. coli* (Buchanan and Sowell, 1982; Li *et al.*, 2014). The loss of DacA results in aberrant cell morphologies, and additional deletions of at least two other PG carboxypeptidases worsen these morphological defects (Nelson and Young, 2000; Ghosh and Young, 2003). DacD (also called PBP6b) was identified as an acidic PG carboxypeptidase whose expression and activity increased at acidic pH (Peters *et al.*, 2016). DacC (also called PBP6a) was recently shown to be involved in overcoming severe OM assembly defects by increasing the number of 3-3 crosslinks of PG (More *et al.*, 2019). DacD and DacC are expressed in mid-exponential growth and stationary phases, respectively (Buchanan and Sowell, 1982; Baquero *et al.*, 1996). DacA, DacC, and DacD are suggested to be membrane-anchored via their C-terminal amphiphilic α-helix domains, and this domain is important for the morphological functioning of PBP5 (Nelson *et al.*, 2002), its localization at the cell division site (Nelson *et al.*, 2002; Potluri *et al.*, 2010), and β-lactam resistance (Park *et al.*, 2022).

In this study, we investigate the coordinated and distinct functions of the PG carboxypeptidases DacC and DacA in adaptation to alkaline and salt stresses and as well as cell shape maintenance. We found that DacC is an alkaline PG carboxypeptidase whose protein stability and enzyme activity are significantly increased at high pH. Both DacC and DacA were necessary for overcoming high pH stress, whereas only DacA was necessary for adaptation to salt stress. Both DacC and DacA plays important roles in maintaining cell shape, but their roles are distinct. Notably, all these functions of DacC and DacA were not associated with LD-transpeptidases, but with DD-transpeptidases (PBPs). We showed that the C-terminal domain-dependent interactions between PG carboxypeptidases and PBPs are required for the maintenance of cell shape and overcoming alkaline and salt stress. Therefore, our study suggests a model whereby PG carboxypeptidases and PBPs cooperate for adaptation to stress and cell shape maintenance.

## Results

### DacC activity is required for adaptation to alkaline stress

To analyze the role of PG carboxypeptidases in overcoming alkaline stress, we examined the alkaline sensitivity of the PG carboxypeptidase-deleted mutant strains. The *dacC* mutant exhibited strong sensitivity to high pH, whereas the growth of other mutant cells was comparable to that of the wild-type (WT) cells under high pH conditions (Figure 1A). This phenotype was complemented by ectopic expression of DacC under the pBAD plasmid with an arabinose-inducible promoter (Figure 1B) but not by the expression of a DacC(S66G) mutant protein, in which the serine residue of the active site is substituted with glycine (Peters *et al.*, 2016). These results indicate that PG carboxypeptidase activity of DacC is necessary for the adaptation to alkaline stress. We examined whether the overexpression of other PG carboxypeptidases could complement the phenotype of the *dacC* mutant. In the presence of 1% arabinose, DacC fully complemented the phenotype of the *dacC* mutant, whereas other proteins hardly complemented it (Figure 1C), suggesting that DacC has a distinct role in overcoming alkaline stress.

**Figure 1.**
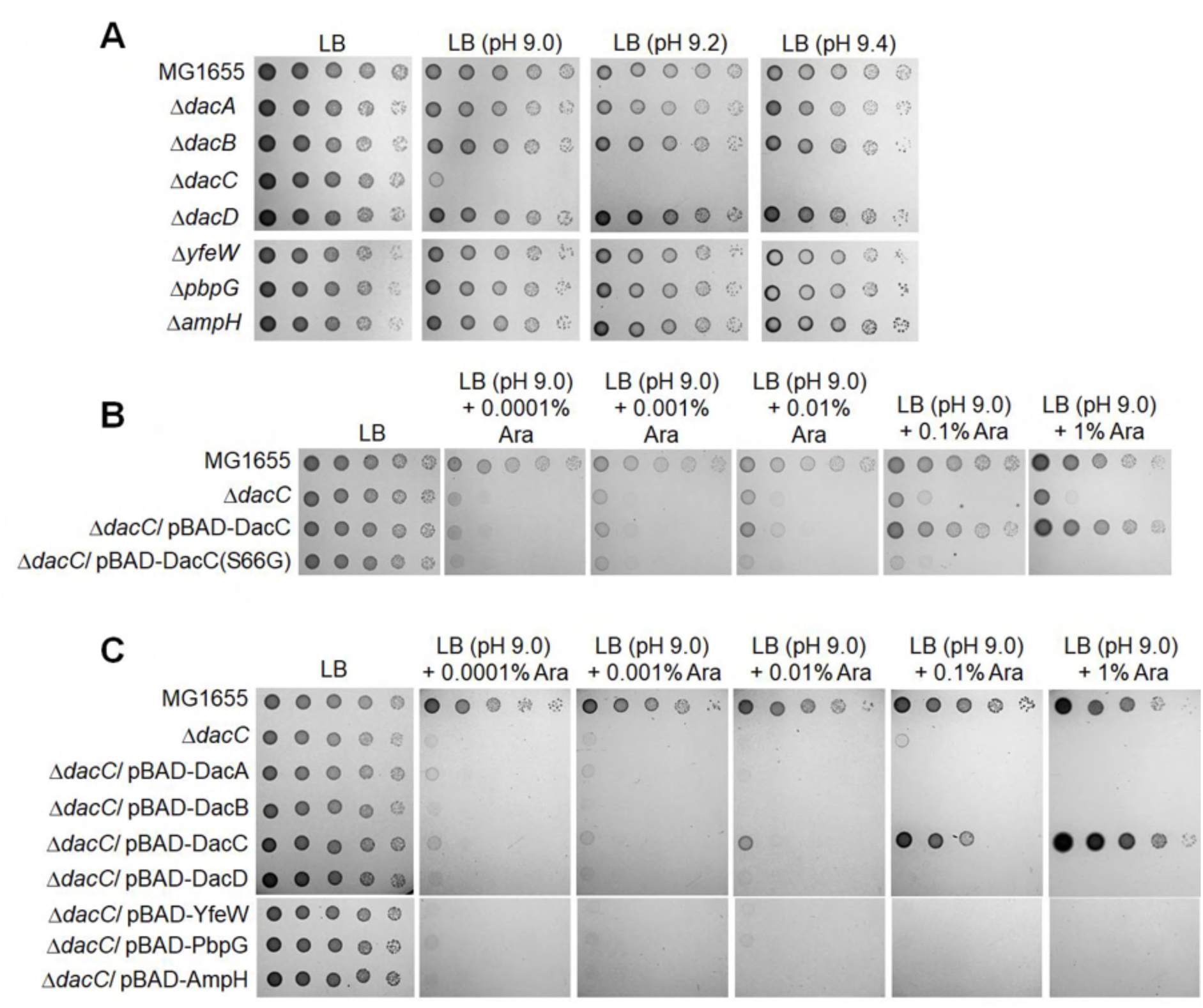
The activity of DacC is required for growth under alkaline stress conditions. (**A**) Sensitivity of the *dacC* mutant to alkaline pH. The wild-type and indicated mutant cells were serially diluted from 10^8^ to 10^4^ cells/ml in 10-fold steps and spotted onto an LB plate or LB plates at indicated pH. (**B**) Complementation of alkaline sensitivity of the *dacC* mutant. The cells of the indicated strains were serially diluted from 10^8^ to 10^4^ cells/ml in 10-fold steps and spotted onto an LB plate and LB plates at pH 9.0 containing the indicated concentrations of arabinose. (**C**) Complementation of alkaline sensitivity of the *dacC* mutant by other PG carboxypeptidases. The cells of the indicated strains were serially diluted from 10^8^ to 10^4^ cells/ml in 10-fold steps and spotted onto an LB plate and LB plates at pH 9.0 containing the indicated concentrations of arabinose.

### Increase in enzyme activity and protein stability of DacC under alkaline stress

A systematic analysis of the protein levels in *E. coli* showed that the protein level of DacC was significantly lower than that of other PG carboxypeptidases under normal growth conditions (Li *et al.*, 2014). However, in this study, only the growth of the *dacC* mutant was inhibited under alkaline stress and this phenotype was complemented only by the expression of DacC (Figure 1). To understand the distinct role of DacC under alkaline stress, we examined enzyme activity and expression levels of DacC under alkaline stress. A previous study also prompted us to investigate this point, showing that the PG carboxypeptidase activity of DacD was higher at low pH than at neutral pH, and its protein level increased at low pH (Peters *et al.*, 2016). An *in vitro* experiment using a bacterial cell wall analog showed that the enzymatic activity of DacC was more than doubled at high pH than at neutral pH (Figure 2A). We examined the mRNA level of the *dacC* gene at high pH. The transcript level of *dacC* did not change under alkaline stress compared to that under normal conditions (Figure 2B). The protein level of DacC was also examined under high pH conditions. The protein level of DacC increased by more than 3-fold under alkaline stress conditions than under normal conditions (Figure 2C and D). Additionally, we checked the protein levels of DacC paralogs, DacA and DacD. The protein level of DacA did not chang under alkaline stress conditions, whereas DacD was strongly diminished under alkaline stress conditions (Figure 2C and D). Therefore, these results indicate that DacC and DacD are PG carboxypeptidases that are specialized for growth at alkaline and acidic pH, respectively. To understand why the protein level of DacC increased under alkaline stress conditions, we estimated the protein stability of DacC. The half-life of DacC was prolonged by more than twice under alkaline stress (approximately 66 min) than under normal conditions (approximately 29 min), indicating that the increased protein level of DacC under alkaline stress is mediated by the decreased degradation rate of DacC.

**Figure 2.**
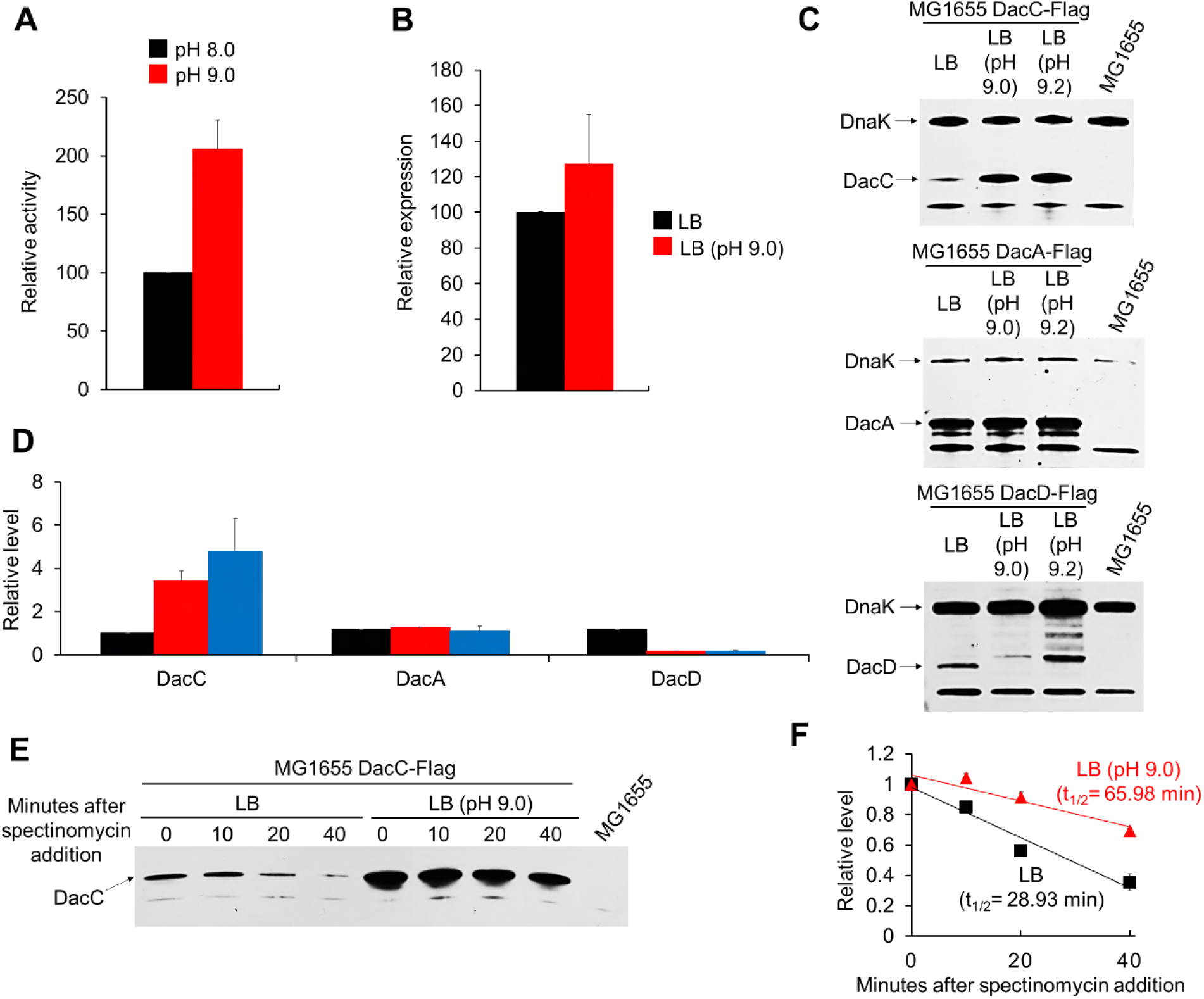
The enzyme activity and protein stability of DacC are enhanced under alkaline stress conditions. (**A**) Increased enzymatic activity of DacC at an alkaline pH. The enzymatic activity of purified DacC was measured using the bacterial cell wall analog, diacetyl-L-Lys-D-Ala-D-Ala (AcLAA). Purified protein (1 μM) was incubated at 37°C for 60 min with 50 mM Tris-HCl (pH 8.0 or 9.0) containing 1 mM AcLAA. The amount of D-Ala released by DacC was measured using horseradish peroxidase and Amplex Red. (**B**) Relative transcript levels of the *dacC* gene in LB medium (black bar) and LB medium at pH 9.0 (red bar). Total mRNA was extracted from MG1655 cells grown in the indicated media to the early exponential phase (OD_600nm_ = 0.4). Data were obtained from three independent experiments. mRNA levels were normalized to the concentration of 16S rRNA. (**C**) Protein levels of DacC, DacA, and DacD in LB medium or LB media at the indicated pH. Western blot analysis with anti-Flag and anti-DnaK antibodies was performed using MG1655 DacC-Flag, DacA-Flag, or DacD-Flag cells grown in the indicated media to the early exponential phase (OD_600nm_ = 0.4). DnaK was used as the loading control. (**D**) Quantification of the protein levels in (**C**) plotted as relative levels: black bars, LB; red bars, LB (pH 9.0); blue bars, LB (pH 9.2). Error bars represent the standard deviation from triplicate measurements. (**E**) *In vivo* degradation assay of DacC. MG1655 DacC-Flag cells were grown in LB medium or LB medium at pH 9.0 to the early exponential phase (OD_600nm_ = 0.4). Spectinomycin was added to each culture to a final concentration of 200 μg/ml to block protein synthesis, and 6 × 10^7^ cells were harvested at the indicated time points for western blot analysis. Each experiment was performed in triplicate and the representative image is shown. (**F**) The normalized signal at the time of spectinomycin addition was set as 1 and the change in signal intensity at each time point was plotted for each medium: black, LB; red, LB (pH 9.0). Error bars represent the standard deviation from the triplicate measurements.

### DacA is necessary for growth under alkaline and salt stresses

Because the protein level of DacA was consistently maintained under alkaline stress (Figure 2C), its role in adaptation to alkaline stress was examined. Although the *dacA* single mutant showed normal growth under alkaline stress (Figure 1A), additional deletion of DacA in the *dacC* mutant worsened the growth defect of the *dacC* mutant under alkaline stress (Figure 3A).

**Figure 3.**
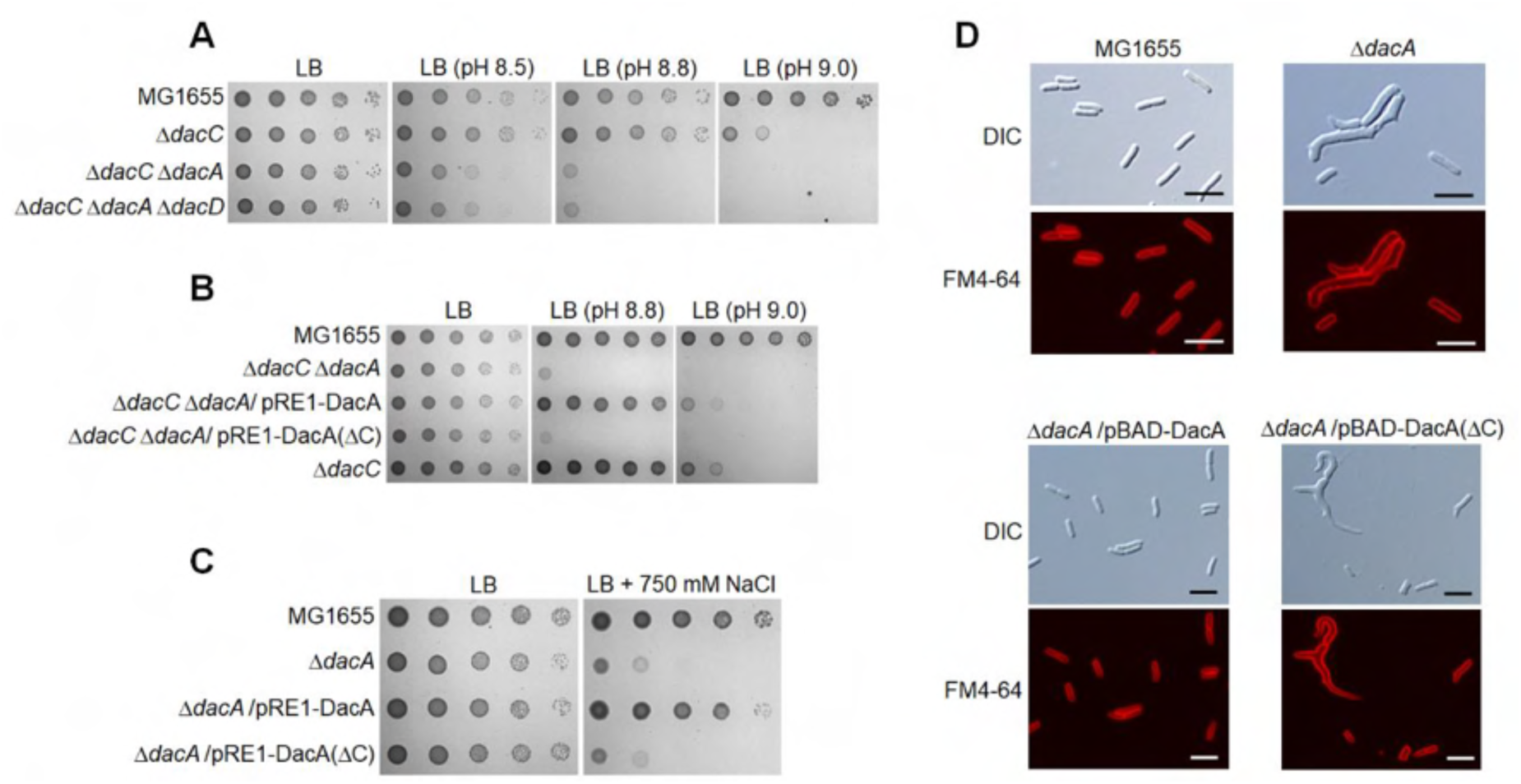
Effects of DacA depletion on morphological maintenance and adaptation to alkaline and salt stresses. (**A**) Increased sensitivity of the *dacC* mutant to alkaline pH by additional deletion of the *dacA* gene. The wild-type and indicated mutant cells were serially diluted from 10^8^ to 10^4^ cells/ml in 10-fold steps and spotted onto an LB plate or LB plates at indicated pH. (**B**) Complementation of alkaline sensitivity of the *dacC dacA* double mutant. The cells of the indicated strains were serially diluted from 10^8^ to 10^4^ cells/ml in 10-fold steps and spotted onto an LB plate and LB plates at the indicated pH. (**C**) Salt sensitivity of the *dacA* mutant. The wild-type and indicated mutant cells were serially diluted from 10^8^ to 10^4^ cells/ml in 10-fold steps and spotted onto an LB plate or an LB plate containing 750 mM NaCl. (**D**) Morphological defects of the *dacA* mutant in LB medium. The indicated cells grown in LB medium containing 0.2% arabinose were stained with FM4-64 (red), and then spotted on a 1% agarose pad. Scale bars, 5 μm.

The additional deletion of DacD in the *dacC dacA* double mutant did not affect bacterial growth under alkaline stress. This phenotype, caused by the additional deletion of DacA in the *dacC* mutant, was fully complemented by the ectopic expression of DacA (Figure 3B). These results indicate that DacA plays an accessory role in adapting to alkaline stress. The *dacA* mutant also exhibited salt sensitivity, which was fully complemented by the ectopic expression of DacA (Figure 3C). This phenotype was not detected in other PG carboxypeptidase mutants, including the *dacC* mutant (Park *et al*., 2022), and additional deletions of DacC and DacD did not increase salt sensitivity of the *dacA* mutant (Figure 3–figure supplement 1), indicating that only DacA was associated with this phenotype. These data suggest that DacA is required to adapt to alkaline and salt stresses.

### Inactivation of DacA and DacC leads to distinct shape abnormalities

DacA has been known to be important for morphological maintenance (Nelson and Young, 2000; Nelson and Young, 2001; Nelson *et al.*, 2002). Cells with an abnormal morphology, such as branched or filamentous shapes, were detected in the *dacA* mutant under normal growth conditions (Figure 3D). These morphological defects were fully complemented by the ectopic expression of DacA. The *dacC* mutant did not exhibit any morphological defects under normal growth conditions (Figure 4A). Next, we examined the effect of alkaline stress on the morphology of the *dacC* and *dacA* mutants. Notably, under alkaline stress, the *dacC* mutant exhibited a spherical shape, whereas the *dacA* mutant exhibited a filamentous shape (Figure 4A). These morphological phenotypes were complemented by the ectopic expression of each protein (Figure 4B and C). It is noteworthy that cells with abnormal morphology were observed in the *dacA* mutant at neutral pH but not at the alkaline pH range of 8.6 and 8.8 (Figure 4A). These results indicate that both of DacA and DacC are necessary for shape maintenance, but their roles in morphology are distinct.

**Figure 4.**
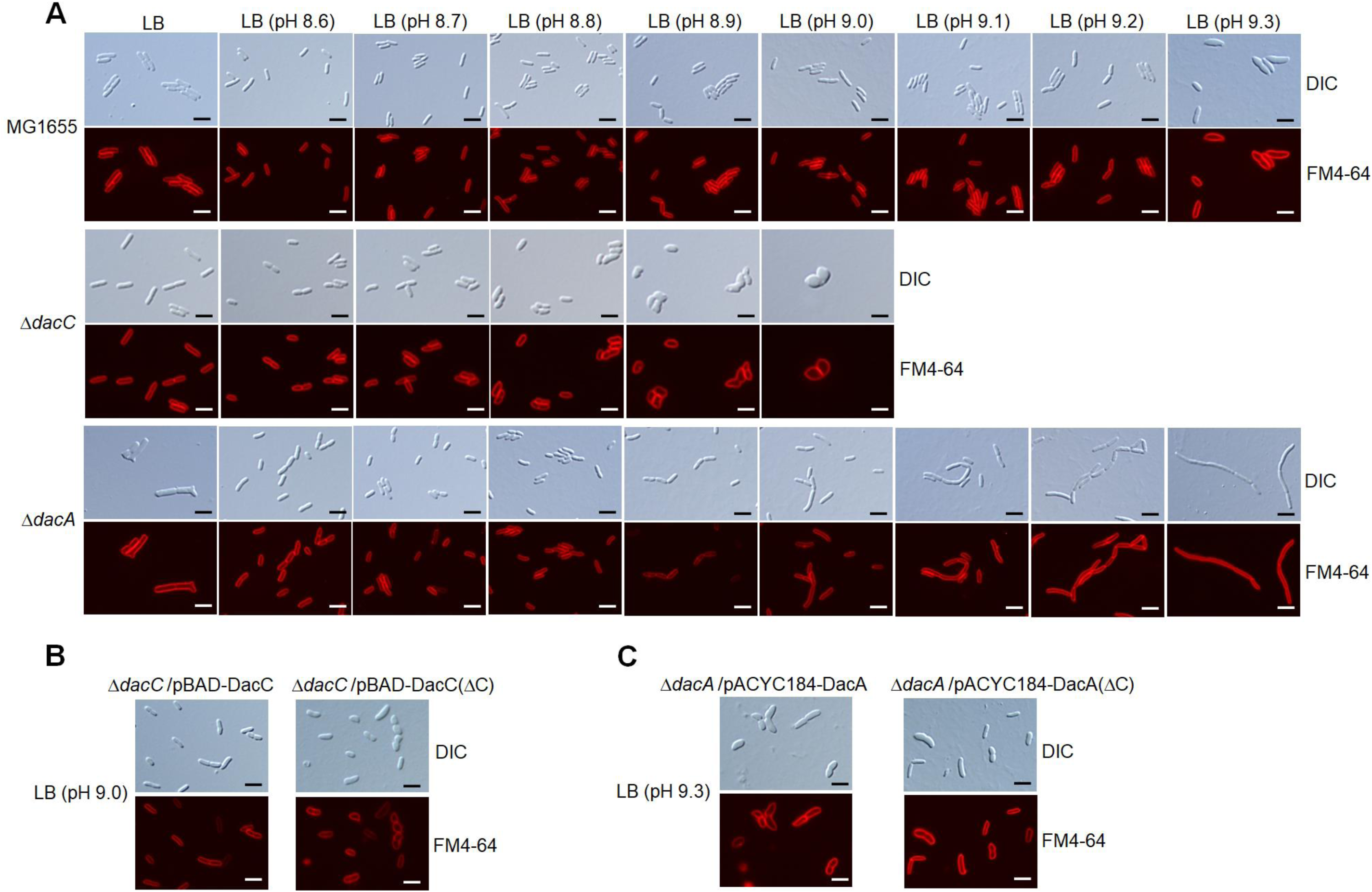
Distinct morphological changes of *dacC* and *dacA* mutants under alkaline stress conditions. (**A**) Cell shapes of *dacC* and *dacA* mutants under alkaline stress conditions. The indicated cells grown in LB medium or LB media at the indicated pH were stained with FM4-64 (red), and then spotted on a 1% agarose pad. Scale bars, 5 μm. (**B**) Complementation of spherical shape of the *dacC* mutant under alkaline stress conditions. The indicated cells grown in LB medium at pH 9.0 containing 0.2% arabinose were stained with FM4-64 (red), and then spotted on a 1% agarose pad. Scale bars, 5 μm. (**C**) Complementation of filamentous shape of the *dacA* mutant under alkaline stress conditions. The indicated cells grown in LB medium at pH 9.3 were stained with FM4-64 (red), and then spotted on a 1% agarose pad. Scale bars, 5 μm.

### The roles of DacC and DacA in stress adaptation and morphology maintenance are not associated with LD-transpeptidases

After the removal of the fifth D-Ala by PG carboxypeptidases, PG stem peptides can be covalently attached to Lpp lipoproteins or adjacent PG peptides by LD-transpeptidases (Hugonnet *et al.*, 2016). The attachment of Lpp lipoproteins is catalyzed by LdtABC (Hugonnet *et al.*, 2016), whereas the formation of 3-3 crosslinks is catalyzed by LdtDE (Bahadur *et al.*, 2021; Winkle *et al.*, 2021). Thus, we examined the relationship between LD-transpeptidases and roles of DacA and DacC identified in this study. Cells defective in all LD-transpeptidases showed normal growth under alkaline stress, comparable to that of the WT strain (Figure 5A). Additionally, the deletion of DacC in the Δ*ldtABCDE* strain caused sensitivity to alkaline stress up to the same level as that of the *dacC* mutant (Figure 5A), indicating that the alkaline sensitivity of the *dacC* mutant is not associated with LD-transpeptidases. Likewise, the salt sensitivity of the *dacA* mutant was not associated with LD-transpeptidases (Figure 5B).

**Figure 5.**
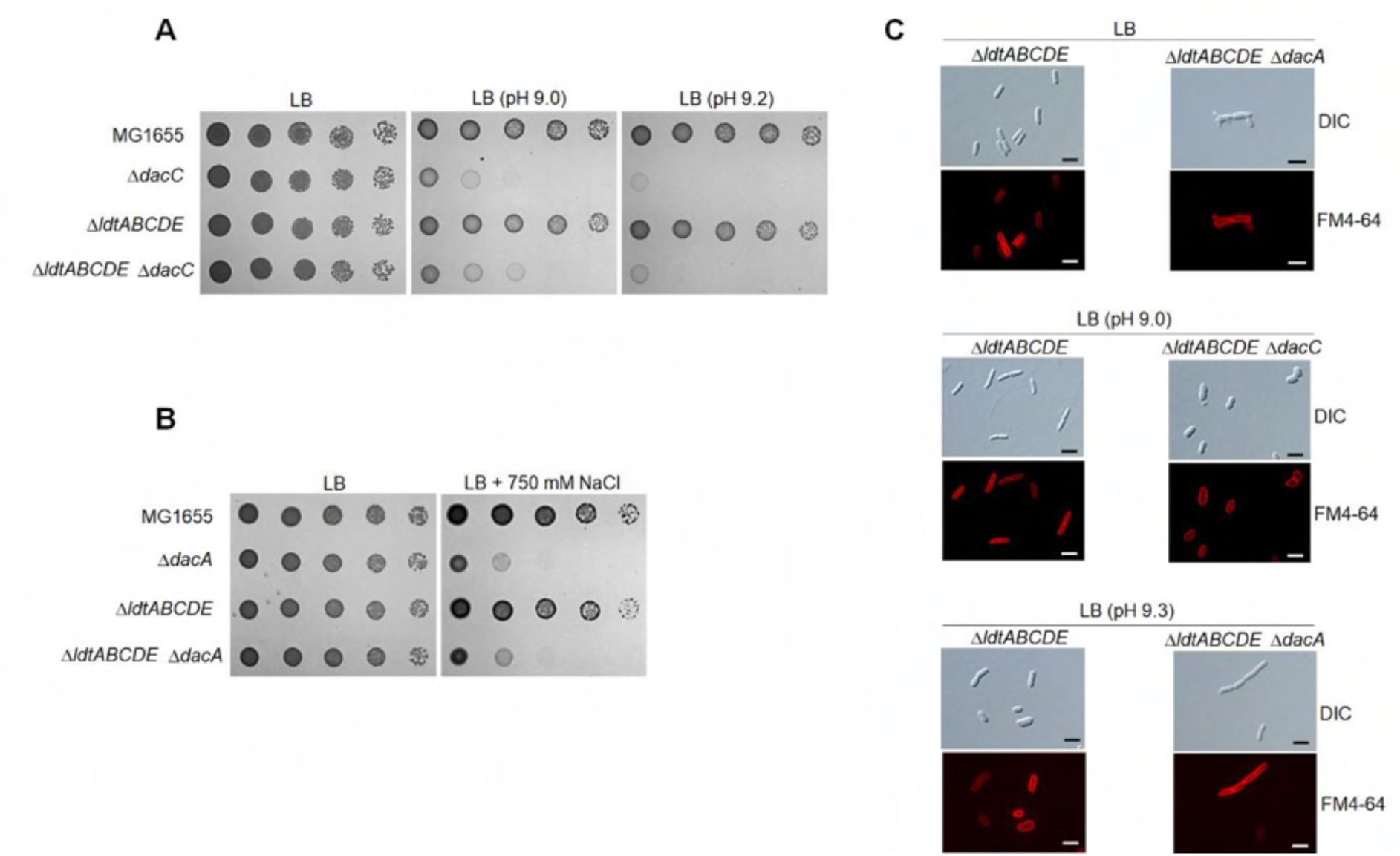
Identified phenotypes of the *dacC* and *dacA* mutants are independent of LD-transpeptidases. (**A**) LD-Transpeptidase-independent alkaline sensitivity of the *dacC* mutant. The wild-type and indicated mutant cells were serially diluted from 10^8^ to 10^4^ cells/ml in 10-fold steps and spotted onto an LB plate or LB plates at indicated pH. (**B**) LD-Transpeptidase-independent salt sensitivity of the *dacA* mutant. The wild-type and indicated mutant cells were serially diluted from 10^8^ to 10^4^ cells/ml in 10-fold steps and spotted onto an LB plate or an LB plate containing 750 mM NaCl. (**C**) LD-Transpeptidase-independent morphological changes of the *dacC* and *dacA* mutants. The indicated cells grown in LB medium or LB media at the indicated pH were stained with FM4-64 (red), and then spotted on a 1% agarose pad. Scale bars, 5 μm.

The *dacA* mutant showed an abnormal shape in neutral pH lysogeny broth (LB) medium and a filamentous shape in alkaline LB medium (Figure 3D, Figure 4A), whereas the *dacC* mutant showed a spherical shape in alkaline LB medium (Figure 4A). However, these morphological abnormalities were not observed in the Δ*ldtABCDE* strain, and the additional deletion of DacA and DacC in the Δ*ldtABCDE* strain phenocopied the morphological defects of the *dacA* and *dacC* mutants, respectively (Figure 5C). Collectively, these results demonstrate that the physiological roles of DacA and DacC in stress adaptation and morphology are not mediated by LD-transpeptidase-related functions.

### C-terminal membrane-anchoring domains of DacC and DacA are indispensable for their most cellular functions, but not their enzymatic activities

DacC has a membrane-anchoring domain at its C-terminus, and a lack of this domain produces a soluble protein (van der Linden *et al.*, 1992; Harris *et al.*, 2002). We constructed a DacC(ΔC) mutant protein defective in the membrane-anchoring domain to investigate the role of this domain in adaptation to alkaline stress. Although the PG carboxypeptidase activity of DacC(ΔC) was significantly higher than that of WT DacC, regardless of pH (Figure 6A), DacC(ΔC) hardly complemented the alkaline sensitivity of the *dacC* mutant (Figure 6B). DacC(ΔC) did not complement morphological changes in the *dacC* mutant under alkaline stress (Figure 4B). To confirm the requirement of the C-terminal domain of DacC in its cellular functions, we constructed a strain defective for the C-terminal domain of chromosomal DacC, MG1655 DacC(ΔC)-Flag. Although the expression level of DacC(ΔC) was comparable to that of DacC (Figure 7A), MG1655 DacC(ΔC)-Flag cells were highly sensitive to alkaline stress (Figure 7B) and exhibited morphological changes under alkaline stress (Figure 7C), similar to the *dacC* mutant. These results imply that retaining only the PG carboxypeptidase activity of DacC is not sufficient for its intracellular role and that another factor is required for its function.

**Figure 6.**
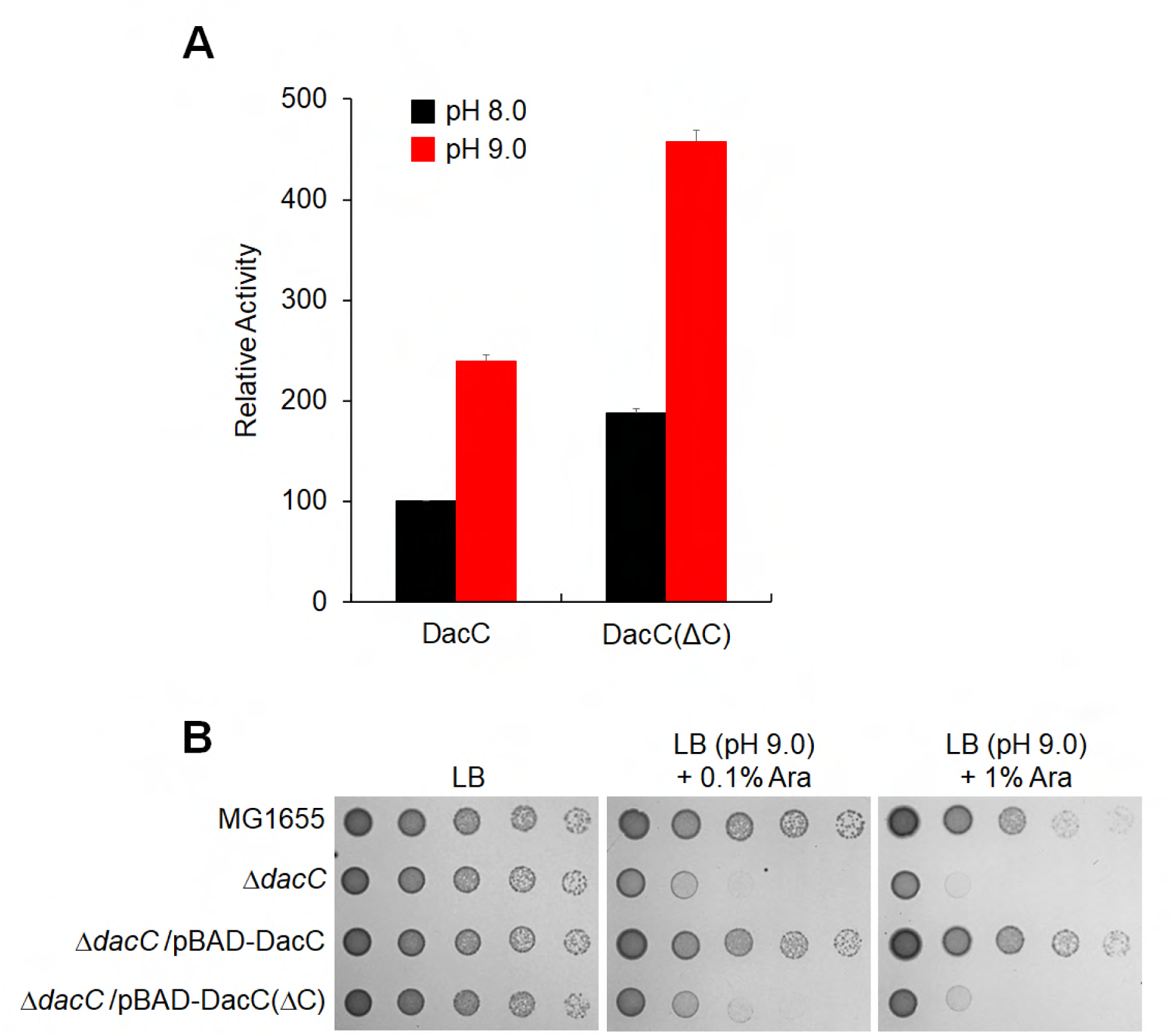
The C-terminal domain of DacC is necessary for adaptation to alkaline stress, but not its PG carboxypeptidase activity. (**A**) Enhanced PG carboxypeptidase activity of DacC(ΔC). The enzymatic activities of purified DacC and DacC(ΔC) were measured using the bacterial cell wall analog, diacetyl-L-Lys-D-Ala-D-Ala (AcLAA). Purified proteins (1 μM) were incubated at 37°C with 50 mM Tris-HCl (pH 8.0 or 9.0) containing 1 mM AcLAA. The amount of released D-Ala was estimated using and Amplex Red and horseradish peroxidase at 563 nm. (**B**) Requirement of the C-terminal domain of DacC to adapt to alkaline stress. The wild-type and indicated mutant cells were serially diluted from 10^8^ to 10^4^ cells/ml in 10-fold steps and spotted onto an LB plate or LB plate at pH 9.0 containing 0.1% or 1% arabinose.

**Figure 7.**
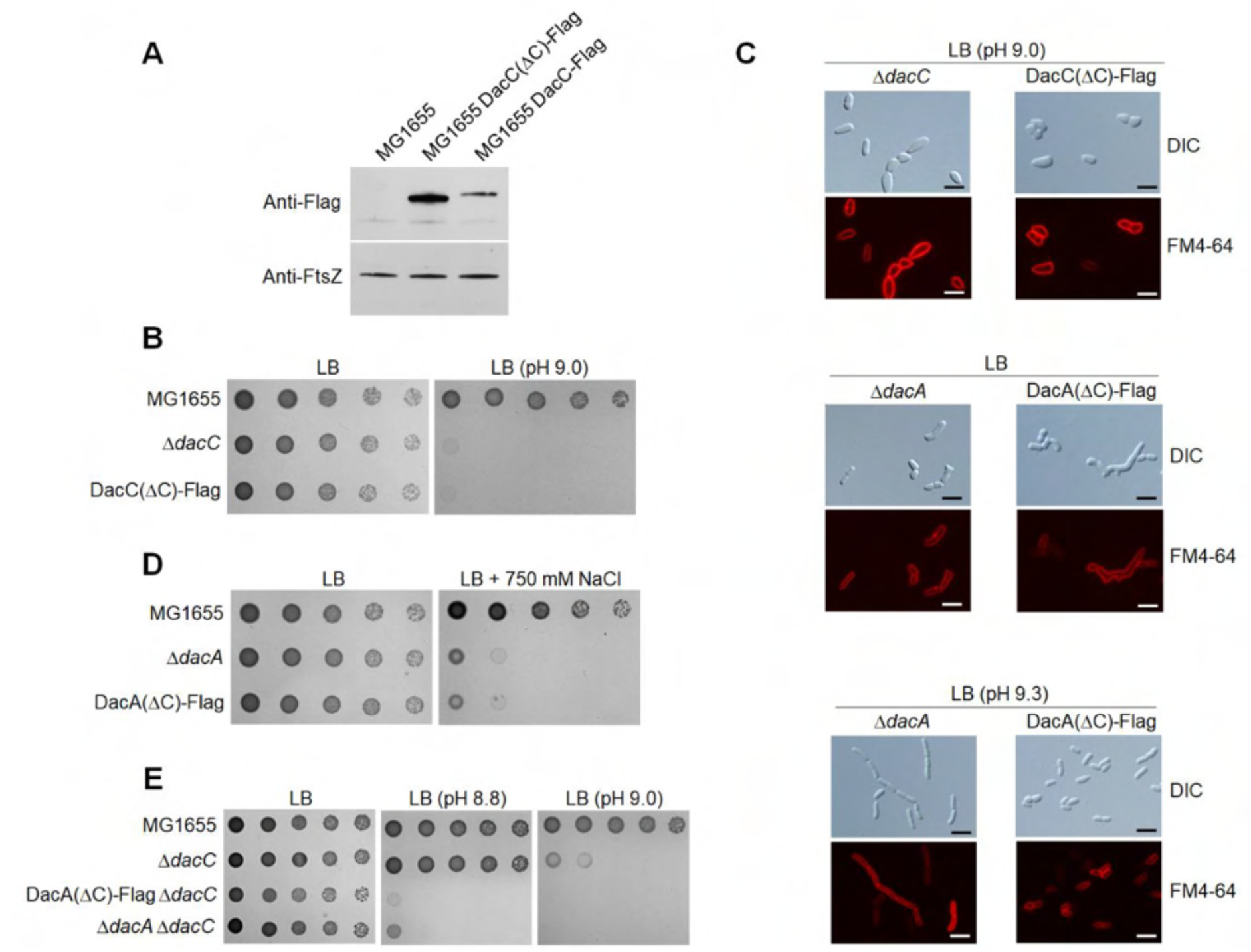
Phenotypes of the MG1655 DacC(ΔC)-Flag and MG1655 DacA(ΔC)-Flag strains. (A) Intracellular protein levels of DacC and DacC(ΔC). Western blot analysis with anti-Flag and anti-FtsZ antibodies was performed using 5 x 10^7^ cells of indicated strains grown in LB medium to the early exponential phase (OD_600nm_ = 0.4). FtsZ was used as the loading control. (B) Sensitivity of the MG1655 DacC(ΔC)-Flag strain to alkaline pH. The wild-type and indicated mutant cells were serially diluted from 10^8^ to 10^4^ cells/ml in 10-fold steps and spotted onto an LB plate or LB plate at pH 9.0. (**C**) Morphological changes of the MG1655 DacC(ΔC)-Flag and MG1655 DacA(ΔC)-Flag strains. The indicated cells grown in LB medium or LB media at the indicated pH were stained with FM4-64 (red), and then spotted on a 1% agarose pad. Scale bars, 5 μm. (**D**) Salt sensitivity of the MG1655 DacA(ΔC)-Flag strain. The wild-type and indicated mutant cells were serially diluted from 10^8^ to 10^4^ cells/ml in 10-fold steps and spotted onto an LB plate or LB plate containing 750 mM NaCl. (**E**) Phenotype of the MG1655 DacA(ΔC)-Flag Δ*dacC* strain under alkaline stress conditions. The wild-type and indicated mutant cells were serially diluted from 10^8^ to 10^4^ cells/ml in 10-fold steps and spotted onto an LB plate or LB plates at the indicated pH.

The role of the C-terminal membrane-anchoring domain of DacA was also analyzed. A previous report showed that PG carboxypeptidase activity of DacA(ΔC) was significantly higher than that of WT DacA (Park *et al.*, 2022). DacA(ΔC) did not complement the alkaline and salt sensitivities of the *dacA* mutant or morphological defects in LB medium (Figure 3). However, DacA(ΔC) fully complemented the filamentous morphology of the *dacA* mutant under alkaline stress (Figure 4C). Additionally, these results were confirmed by using a strain defective for the C-terminal domain of chromosomal DacA, MG1655 DacA(ΔC)-Flag. Although the expression level of DacA(ΔC) was comparable to that of DacA (Park *et al.*, 2022), MG1655 DacA(ΔC)-Flag cells were highly sensitive to alkaline and salt stresses (Figure 7D and E) and exhibited morphological defects in LB medium (Figure 7C). However, MG1655 DacA(ΔC)-Flag cells did not exhibit filamentous morphology under alkaline stress (Figure 7C). Taken together, these results demonstrate that the PG carboxypeptidase activities of DacC and DacA are not solely sufficient for most intracellular roles and that other factors are required for their functions.

### DacC and DacA directly interact with PBPs, mostly in a C-terminal domain-dependent manner

In a previous study, we demonstrated that DacA directly interacts with PBPs in a C-terminal domain-dependent manner (Park *et al*., 2022). To determine whether these interactions are also detected between DacC and PBPs, we performed pull-down experiments using DacC-Flag and DacC(ΔC)-Flag strains expressing His-tagged or non-tagged PBPs. Despite similar intracellular expression levels of DacC, DacC was pulled down only by His-tagged PBPs and not non-tagged PBPs (Figure 8A), indicating physical interactions between DacC and PBPs. When the same experiment was performed using the DacC(ΔC)-Flag strain, DacC(ΔC) was not pulled down by His-tagged PBPs, except for PBP2 (Figure 8A), despite the higher expression levels of DacC(ΔC) than those of DacC (Figure 8A) and similar levels of His-tagged PBPs eluted from DacC-Flag and DacC(ΔC)-Flag strains (Figure 8–figure supplement 1). These results indicate that the C-terminal domain of DacC is required for physical interactions with PBPs, except for PBP2. Under the same experimental conditions, DacA also interacted with all PBPs, and its C-terminal domain was required for physical interactions with all PBPs, including PBP2 (Figure 8B, Figure 8–figure supplement 2). Collectively, these data show that both DacC and DacA physically interact with all PBPs and that their C-terminal domains are necessary for most interactions with PBPs.

**Figure 8.**
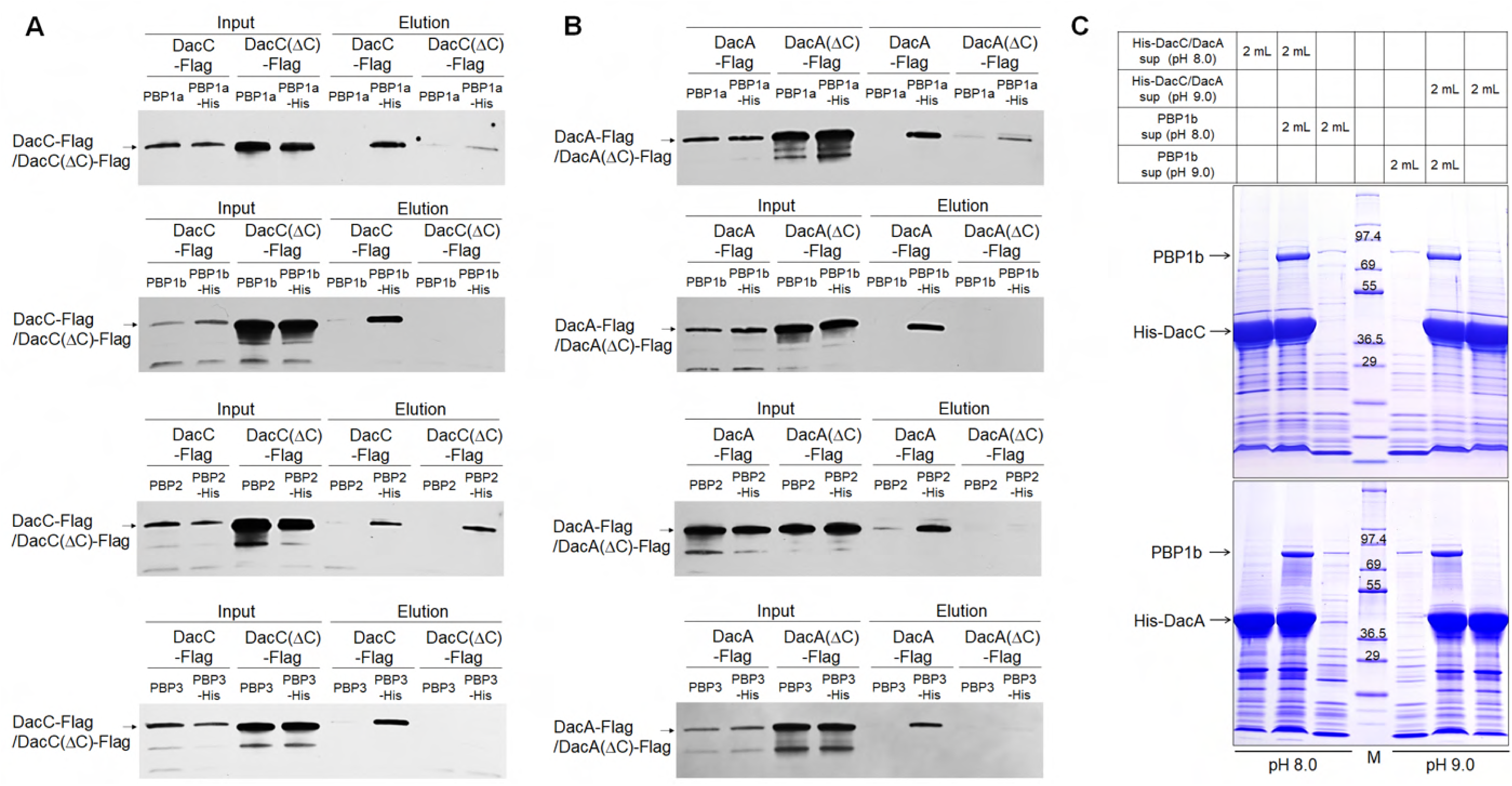
Physical interactions between PBPs and DacC or DacA. (**A**) The C-terminal domain-dependent or -independent interactions of DacC with PBPs. The supernatant of MG1655 DacC-Flag or DacC(ΔC)-Flag cells harboring the pBAD-based plasmids expressing C-terminal His-tagged PBPs or non-tagged PBPs was loaded onto Talon metal affinity resin. After pull-down experiments, the amount of input (Input) and output (Elution) DacC or DacC(ΔC) was measured by western blot using monoclonal antibody against Flag-tag. (**B**) The C-terminal domain-dependent interactions of DacA with PBPs. The supernatant of MG1655 DacA-Flag or DacA(ΔC)-Flag cells harboring the pBAD-based plasmids expressing C-terminal His-tagged PBPs or non-tagged PBPs was loaded onto Talon metal affinity resin. After pull-down experiments, the amount of input (Input) and output (Elution) DacA or DacA(ΔC) was measured by western blot using monoclonal antibody against Flag-tag. (**C**) The physical interaction between PBP1b and DacC or DacA under alkaline stress conditions. The supernatant of ER2566 cells harboring the pET24a plasmid expressing PBP1b was mixed with the supernatant of ER2566 cells harboring pET28a plasmid expressing His-tagged DacC or His-tagged DacA at pH 8.0 or 9.0. After pull-down experiments, eluted proteins were separated on 4–20% gradient Tris–glycine polyacrylamide gels and visualized by staining with Coomassie brilliant blue R. Lane M indicates EzWay Protein Blue MW Marker (KOMA Biotech).

Because both DacC and DacA are necessary for adaptation to alkaline stress (Figures 1 and 3), we checked whether the physical interaction between PBP and DacC or DacA is retained under alkaline stress conditions. This question was addressed by pull-down experiments using overexpressed PBP1b and His-tagged DacC or DacA. Significant amounts of PBP1b were pulled down by His-tagged DacC and DacA at both pH 8.0 and 9.0 (Figure 8C). PBP1b was not pulled down by His-tagged DacC(ΔC) and DacA(ΔC) at both pH 8.0 and 9.0 (Figure 8–figure supplement 3), although the levels of purified His-tagged DacC(ΔC) and DacA(ΔC) were higher than those of purified His-tagged DacC and DacA, respectively (Figure 8C, Figure 8–figure supplement 3). These results demonstrate the C-terminal domain-dependent interactions of DacC and DacA with PBP1b under alkaline stress conditions.

### aPBP and bPBP are required for stress adaptation and shape maintenance, respectively

If the physical interactions of DacC and DacA with PBPs are important for their physiological roles, strains defective in PBPs could phenocopy the *dacC* or *dacA* mutants. Because PBP2 and PBP3 are essential (Rohs and Bernhardt, 2021), we measured the phenotype of the strain defective for PBP1a (encoded by the *mrcA* gene) or PBP1b (encoded by the *mrcB* gene). Notably, the *mrcB* mutant showed strong sensitivity to alkaline stress, whereas the *mrcA* mutant showed normal growth under alkaline stress conditions (Figure 9A). Additionally, only the mutant defective in LpoB lipoprotein, an adaptor protein for the function of PBP1b, was also significantly sensitive to alkaline stress (Figure 9A). These phenotypes were complemented by PBP1b or LpoB (Figure 9B), indicating that PBP1b, but not PBP1a, is related to adaptation to alkaline stress. Given that PBP1b is an aPBP with both DD-transpeptidase and glycosyltransferase activities, we examined the activity of PBP1b involved in adaptation to alkaline stress. To address this question, we constructed two mutants, PBP1b(E233Q)— defective in glycosyltransferase activity—and PBP1b(S510A)—defective in DD-transpeptidase activity. Neither PBP1b(E233Q) nor PBP1b(S510A) complemented the phenotype of the *mrcB* mutant (Figure 9C), indicating that both these activities are involved in adaptation to alkaline stress. However, because the DD-transpeptidase activity of PBP1b is known to be dependent on glycosyltransferase activity (Bertsche *et al.*, 2005; Egan *et al.*, 2014; Mueller *et al.*, 2019), we cannot determine whether the result of PBP1b(E233Q) is caused by the loss of glycosyltransferase activity or by the glycosyltransferase defect-mediated loss of the DD-transpeptidase activity. Anyway, our results show that the DD-transpeptidase activity of PBP1b is required for adaptation to alkaline stress. To determine whether PBP2 and PBP3 are associated with adaptation to alkaline stress, we used two antibiotics, mecillinam and aztreonam, which exclusively inhibit PBP2 and PBP3, respectively (Kocaoglu and Carlson, 2015; Mueller *et al.*, 2019). If PBP2 and PBP3 are important for adaptation to alkaline stress, WT MG1655 cells may be more sensitive to mecillinam and aztreonam under alkaline stress than under normal conditions. However, the MG1655 cells were more resistant to mecillinam and aztreonam under alkaline stress than under normal conditions (Figure 9D). These results imply that the function of PBP1b, but not PBP1a, PBP2, or PBP3, is necessary for adaptation to alkaline stress. A previous report has shown that both *mrcA* and *mrcB* mutants are highly sensitive to salt stress (Park *et al.*, 2020), indicating that both PBP1a and PBP1b are required for adaptation to salt stress. Similarly, using mecillinam and aztreonam, we analyzed whether PBP2 and PBP3 are associated with adaptation to salt stress. MG1655 cells were slightly more sensitive to mecillinam and aztreonam under salt stress than under normal conditions (Figure 9D), implying that the functions of PBP2 and PBP3 are not highly associated with adaptation to alkaline stress. Collectively, these results suggest that aPBP, but not bPBP, is necessary for stress adaptation. This conclusion is consistent with a previous report showing that aPBP is involved in the repair of PG defects, but not in morphological maintenance (Vigouroux *et al.*, 2020).

**Figure 9.**
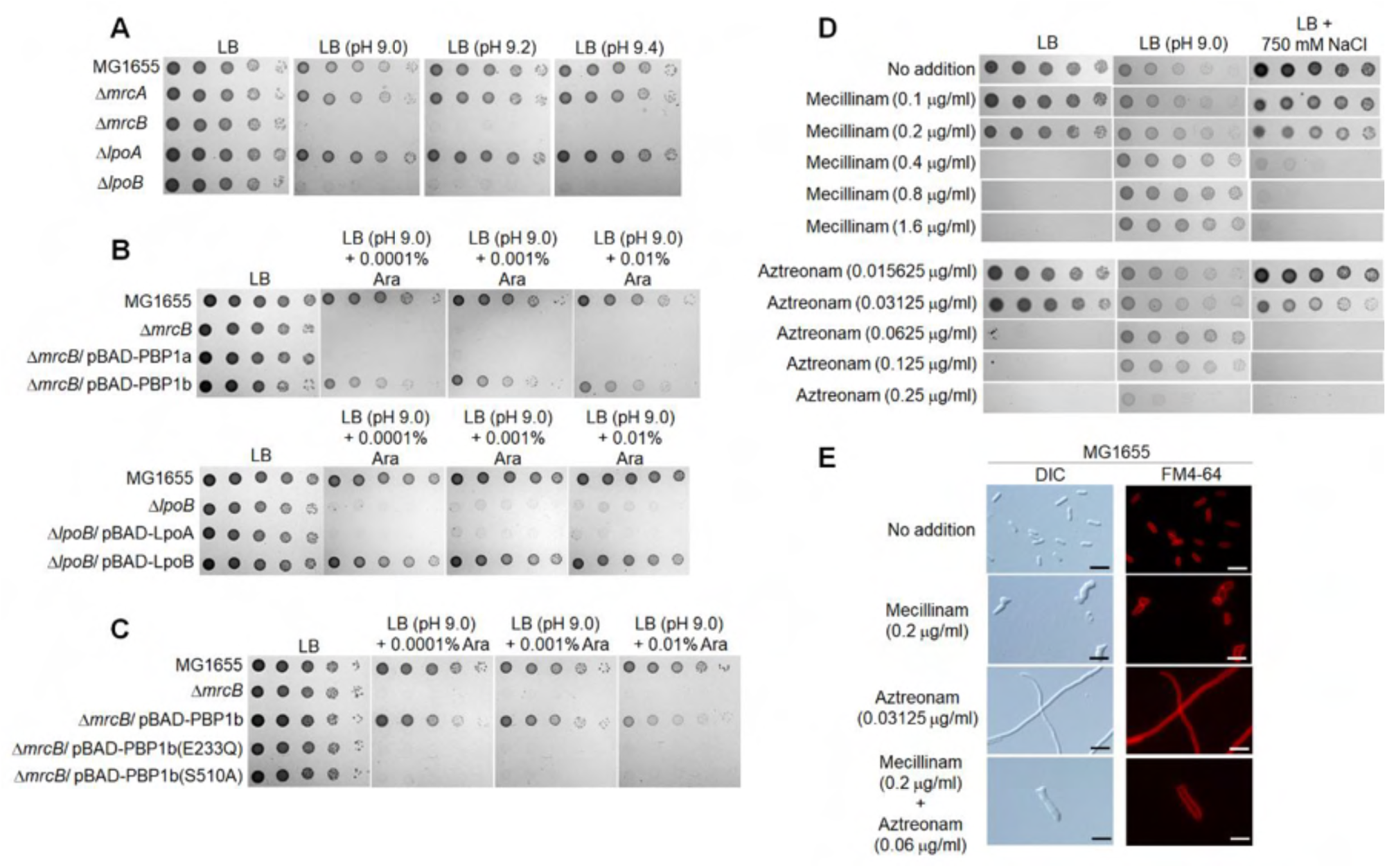
aPBP and bPBP are required for adaptation to alkaline stress and shape maintenance, respectively. (**A**) Sensitivity of *mrcB* and *lpoB* mutants to alkaline stress. The wild-type and indicated mutant cells were serially diluted from 10^8^ to 10^4^ cells/ml in 10-fold steps and spotted onto an LB plate or LB plates at the indicated pH. (**B**) Complementation of alkaline sensitivity of *mrcB* and *lpoB* mutants. The cells of the indicated strains were serially diluted from 10^8^ to 10^4^ cells/ml in 10-fold steps and spotted onto an LB plate and LB plates at pH 9.0 containing the indicated concentrations of arabinose. (**C**) Requirement of the DD-transpeptidase activity of PBP1b to adapt to alkaline stress. The cells of the indicated strains were serially diluted from 10^8^ to 10^4^ cells/ml in 10-fold steps and spotted onto an LB plate and LB plates at pH 9.0 containing the indicated concentrations of arabinose. (**D**) Effect of PBP2 or PBP3 inhibition on adaptation to alkaline or salt stress. The MG1655 cells were serially diluted from 10^8^ to 10^4^ cells/ml in 10-fold steps and spotted onto an LB plate, LB plate at pH 9.0, or LB plate with 750 mM NaCl in the presence of the indicated concentrations of mecillinam or aztreonam. (**E**) Effect of PBP2 or/and PBP3 inhibition on bacterial morphology. The MG1655 cells grown in LB media containing indicated concentrations of mecillinam or/and aztreonam were stained with FM4-64 (red), and then spotted on a 1% agarose pad. Scale bars, 5 μm.

Because the deletion of aPBP does not induce any morphological change (Vigouroux *et al.*, 2020; Rohs and Bernhardt, 2021), the morphological change in the *dacA* or *dacC* mutant may be related to PBP2 and PBP3. However, the shape of the *dacA* mutant in LB medium was different from both the spherical shape caused by inhibition of PBP2 and filamentous shape caused by inhibition of PBP3 (Mueller *et al.*, 2019; Rohs and Bernhardt, 2021). As DacA interacts with both PBP2 and PBP3, the morphological change in the *dacA* mutant may be associated with both PBP2 and PBP3. To address this question, we measured the morphological changes in the presence of mecillinam and aztreonam. MG1655 cells showed spherical or filamentous shapes in the presence of mecillinam or axtreonam, respectively, whereas in the presence of a combination of mecillinam and aztreonam, they showed a branched filamentous shape, which is similar to the shape of the *dacA* mutant (Figure 9E), implying that the morphological change in the *dacA* mutant under normal conditions may be associated with both PBP2 and PBP3. The morphology of the *dacC* mutant under alkaline stress conditions significantly resembled a spherical shape upon inhibition of PBP2 (Figures 4 and 9E). Therefore, these results imply that the morphological change in the *dacA* or *dacC* mutant may be related to bPBP.

## Discussion

PG is an essential bacteria-specific architecture that is necessary for shape maintenance and adaptation to diverse environmental stresses. In this study, we demonstrated that PG carboxypeptidases DacA and DacC are necessary for shape maintenance and adaptation to environmental stresses. DacA and DacC interacted with both aPBP and bPBP, mostly in a C-terminal domain-dependent manner, and these interactions were required for most of their functions in shape maintenance and adaptation to environmental stresses. Notably, these roles of DacA and DacC were not dependent on LD-transpeptidases. These results demonstrate that DacA and DacC play LD-transpeptidase-independent physiological roles in shape maintenance and stress adaptation, which are associated with physical interactions with PBPs.

We found that DacA and DacC play coordinated and distinct roles in stress adaptation and shape maintenance. Both DacC and DacA were required to overcome alkaline stress, whereas only DacA was required to overcome salt stress (Figure 3, Figure 3–figure supplement 1). Both DacA and DacC were necessary for morphological maintenance, but their effects were distinct. Under normal conditions, loss of DacA induced aberrant cell morphology, whereas loss of DacC hardly affected cell morphology (Figure 4). Under alkaline stress conditions, loss of DacA induced a filamentous shape, whereas loss of DacC induced a spherical shape (Figure 4). Basically, these distinct functions of DacA and DacC appear to be based on their protein levels. DacC is an alkaline PG carboxypeptidase, whose stability and activity are significantly increased under alkaline stress conditions (Figure 2). Therefore, in LB medium, only DacA appears to play a predominant role in shape maintenance and stress adaptation. Since DacD was reported to be an acidic PG carboxypeptidase whose stability and activity are increased under acidic stress conditions (Peters *et al.*, 2016), DacD seems to play a minor role under normal conditions. To date, we do not know why DacA and DacC show distinct morphological effects under alkaline stress conditions.

We demonstrated that DacA, DacC, and PBP1b are required to overcome alkaline stress (Figures 3 and 9), whereas PBP1a, PBP2, and PBP3 are not associated with adaptation to alkaline stress (Figure 9). Alkaline stress is known to strongly induce a CpxAR two-component system that senses envelope stress (Danese and Silhavy, 1998; Kumar *et al.*, 2017; Mitchell and Silhavy, 2019), but its molecular mechanism has not been elucidated. As our results imply that alkaline stress might affect PG synthesis or stability, the effect of alkaline stress on PG should be analyzed in further studies.

All PG carboxypeptidase-mediated phenotypes identified in this study were independent of LD-transpeptidases. In the absence of all LD-transpeptidases, all phenotypes of the *dacA* and *dacC* mutants were observed (Figure 5). The C-terminal domain of DacC was dispensable for its enzymatic activity but was required for complementation of all phenotypes of the *dacC* mutant and physical interactions with PBPs, except for PBP2. The C-terminal domain of DacA is dispensable for its enzymatic activity (Park *et al.*, 2022), but was required for complementation of all phenotypes of the *dacA* mutant, except for filamentous morphology under alkaline stress, and all physical interactions with PBPs. These results suggest that many functions of DacA and DacC are not linked to LD-transpeptidase, but are instead associated with DD-transpeptidases, PBPs. At present, we do not know how DacA and DacC affect the functions of PBPs or act cooperatively in PG synthesis with PBPs. We previously reported divergent phenotypes of the *dacA* mutant toward antibiotics, vancomycin resistance and β-lactam sensitivity (Park *et al.*, 2022). The β-lactam sensitivity of the *dacA* mutant was dependent on the C-terminal domain of DacA, whereas vancomycin resistance was independent of its C-terminal domain (Park *et al.*, 2022), like filamentous morphology under alkaline stress in this study. These results indicate that there are cellular processes required for only the enzymatic activity of DacA, and that the cellular functions of DacA are more diverse than expected. Based on the phenotypes of PBP-defective mutants, we predicted which PBPs were associated with the phenotypes of the *dacC* and *dacA* mutants. aPBPs appear to be involved in adaptation to alkaline and salt stresses, whereas bPBPs appear to be involved in shape maintenance (Figure 9). These results are consistent with a previous study showing that aPBPs contribute to PG repair rather than to morphological maintenance (Vigouroux *et al.*, 2020). Further studies are required to elucidate the interactions with PBPs that are linked to the physiological roles of DacA and DacC.

In this study, we identified novel roles for DacA and DacC in shape maintenance and adaptation to alkaline and salt stresses. These roles of DacA and DacC were independent of LD-transpeptidase but were instead associated with physical interactions with PBPs in a C-terminal domain-dependent manner. Additionally, we revealed that DacC is an alkaline PG carboxypeptidase, whose activity and stability are significantly enhanced under alkaline stress conditions, indicating the physiological significance of PG carboxypeptidase redundancy. Therefore, our results provide novel insights and direction for the study of PG carboxypeptidases.

## Materials and Methods

### Bacterial strains, plasmids, and culture conditions

All *E. coli* strains used in this study are listed in Table S1, and all the primer sequences used in this study are presented in Table S2. All cells were cultured in LB medium at 37°C, unless otherwise stated. Ampicillin (100 μg/ml), kanamycin (50 μg/ml), chloramphenicol (5 μg/ml), and tetracycline (10 μg/ml) were added into the culture medium, when necessary. Alkaline LB medium was prepared by adding 50 mM Tris (final concentration), followed by pH adjustment of the medium using 10 N NaOH solution. The final pH of medium was defined as the pH of medium after autoclave sterilization process, due to slight decrease of pH after autoclave process.

All deletion strains were constructed using the plasmid pKD46 expressing λ red recombinase, as previously described (Datsenko and Wanner, 2000), with some modifications. Deletion cassettes for the exchange of the target gene with the FRT sequence containing the kanamycin resistance gene were amplified from the pKD13 plasmid. After polymerase chain reaction (PCR) purification, deletion cassettes were transformed into MG1655 cells harboring the pKD46 plasmid. Transformed cells were spread onto LB medium plates containing kanamycin and plates were incubated at 37°C or 30°C overnight. The deletion strain was confirmed using PCR. To remove the kanamycin resistance gene, the pCP20 plasmid expressing FLP recombinase, which catalyzes recombination between FRT sequences, was transformed into the deletion strain. Removal of the kanamycin resistance gene was also confirmed using PCR. To minimize the impact on the physiology of the deletion strain, the pCP20 plasmid was removed at 37°C, as previously reported (Park *et al*., 2020).

The insertion of a *3× Flag* gene into the chromosomal 3’ region of the target gene was performed using λ red recombinase, as previously reported (Kim *et al.*, 2021; Park *et al.*, 2022). The plasmid pBAD-Flag-FRT-Kan, which contained both the *3× Flag* gene and the FRT-flanked kanamycin resistance gene, was used as a template for PCR amplification using the primer sets listed in Table S2. After PCR amplification, the pBAD-Flag-FRT-Kan plasmid, which was used as a template, was removed by overnight treatment with the restriction enzyme *Dpn*I. The insertion of both the *3× Flag* gene and kanamycin resistance gene was conducted in the MG1655 strain harboring the pKD46 plasmid. After chromosomal fusion of the 3×Flag-tag, the kanamycin resistance gene was removed using FLP recombinase of the pCP20 plasmid, as described above. In the MG1655 DacC(ΔC)-Flag strain, the 3×Flag epitope was inserted at the 378^th^ amino acid (glycine residue) of DacC instead of its C-terminal amino acid (serine residue).

To construct the pBAD-DacC plasmid, the region covering the entire open leading frame of the *dacC* gene was amplified. After PCR purification, the *dacC* gene was inserted into the pBAD24 plasmid digested with EcoRI and XbaI. The insertion was conducted by recombination of overlapping sequences using Infusion cloning (Clontech, USA), as previously reported (Park *et al.*, 2020). The cloning of the *dacC* gene was confirmed by sequence analysis. The plasmid pBAD-DacC(S66G) was constructed by PCR using pBAD-DacC as a template and DpnI-dependent digestion of the template plasmid. The other pBAD-based plasmids were constructed using the same method. The pET24a-based plasmids expressing non-tagged WT proteins and pET28a-based plasmids expressing His-tagged proteins were constructed using similar methods, except for NdeI and BamHI restriction enzymes used for plasmid digestion. pRE1-based plasmids were constructed using similar methods, except for NdeI and XbaI restriction enzymes used for plasmid digestion. Because the gene in the pRE1 plasmid is transcribed by *E. coli* RNA polymerase under the control of a combination of strong λ *P*_L_ promoter and cI ribosome binding site, it is constitutively transcribed in general *E. coli* strains without the cI protein, such as MG1655 (Reddy *et al.*, 1989; Lee *et al.*, 2021). The pACYC184-based plasmids were constructed using similar methods, except for BamHI and EagI restriction enzymes used for plasmid digestion. In this study, the region covering both the promoter region and the entire open leading frame of the target gene was cloned into the pACYC184 plasmid. All the primers used for plasmid construction are listed in Table S2.

### Quantitative real-time PCR

To measure the mRNA level of *dacC* at high pH, total RNA was extracted from the cells cultured in LB medium or LB medium buffered to pH 9.0. Total RNA was extracted when the cells reached an OD_600_ of approximately 0.4. To remove genomic DNA, each RNA sample was incubated with RNase-free DNase I (Promega, USA) at 37°C for 1 h. The remaining RNA was converted into cDNAs using the cDNA EcoDry Premix (Clontech, USA). Quantitative real-time PCR was performed using 10-fold diluted cDNAs as the template. Primers for the detection of *dacC* or 16S rRNA listed in Table S2 were used, and amplification was performed using the SYBR Premix Ex Taq II (Takara, Japan) solution. PCR and simultaneous fluorescence detection were performed using a CFX96 Real-Time System (Bio-Rad, USA). The expression level of *dacC* was calculated as the difference between the threshold cycle of *dacC* and that of the 16S rRNA reference gene.

### Detection of intracellular levels of PG carboxypeptidases

The protein levels of DacA, DacC, and DacD were detected using MG1655 strains with chromosomal insertion of the *3× Flag* gene into the 3’ region of each gene. The cells were cultured in LB medium or LB medium buffered to pH 9.0 or 9.2 and were harvested at an OD_600_ of approximately 0.4. After the addition of sodium dodecyl sulfate sample buffer, the samples were boiled at 100°C for 5 min. Total proteins were separated on 4–20% Tris–glycine polyacrylamide gels (KOMA Biotech, Korea) and were transferred onto PVDF membranes. The protein levels of PG carboxypeptidases and DnaK were determined using monoclonal antibodies against Flag-tag (Santa Cruz Biotechnology, USA) and anti-DnaK (Abcam, USA), respectively, according to the standard procedures. DnaK was used as a loading control. The protein level of DacC(ΔC) was detected using similar procedures, except that FtsZ was used as a loading control.

### Measurement of protein stability of DacC

Protein stability of DacC was measured in LB medium and LB medium buffered to pH 9.0. The MG1655 DacC-Flag strain was cultured to an OD_600_ of approximately 0.4. After addition of spectinomycin (200 μg/ml), 6 × 10^7^ cells were harvested at 10, 20, and 40 min. The protein level of DacC was detected using a monoclonal antibody against the Flag-tag (Santa Cruz Biotechnology), according to the western blot protocol described above.

### Purification of PG carboxypeptidase proteins

ER2566 cells harboring the pET28a-based plasmid expressing His-tagged DacC or DacC(ΔC) were cultured in LB medium at 37°C. When the OD_600_ was approximately 0.5, 1 mM isopropyl-β-D-1-thiogalactopyranoside (IPTG) was added to 200 ml of LB medium and cells were cultured at 16°C overnight. Harvested cells were suspended in 3 ml of binding buffer (50 mM Tris-HCl [pH 8.0], 200 mM NaCl, and 1% sodium n-dodecyl-β-D-maltoside) and disrupted twice using a French press at 10,000 psi. The disrupted cells were separated by centrifugation at 9,000 × *g* for 30 min at 4°C, and the supernatant was loaded into 0.5 ml of a Talon metal affinity resin (Clontech) equilibrated with binding buffer. After washing the resin with 3 ml of binding buffer three times, the bound proteins were eluted using elution buffer (50 mM Tris-HCl [pH 8.0], 150 mM NaCl, and 200 mM imidazole). To remove imidazole, the eluted samples were dialyzed with 3 L dialysis buffer (50 mM Tris-HCl [pH 8.0], 50 mM NaCl). The purified proteins were promptly used to assess the enzymatic activity of PG carboxypeptidases.

### Assessment of the enzymatic activity of PG carboxypeptidases

The enzymatic activity of purified His-tagged DacC or DacC(ΔC) was measured using the bacterial cell wall analog, diacetyl-L-Lys-D-Ala-D-Ala (AcLAA) (Peters *et al.*, 2016). D-Ala released by DacC or DacC(ΔC) was degraded into hydrogen peroxide, pyruvate, and amine group by D-amino acid oxidase (Sigma Aldrich). Hydrogen peroxide was reduced to H_2_O by horseradish peroxidase (Sigma Aldrich) using Amplex Red (Invitrogen, USA) as the electron donor. Then, Amplex Red was converted to resorufin by oxidation, and the amount of resorufin was spectrophotometrically measured at 563 nm. The purified His-tagged DacC or DacC(ΔC) protein (1 μM) was mixed with 50 mM Tris-HCl (pH 8.0 or 9.0) containing 1 mM AcLAA in a final volume of 200 μl. After incubation for 60 min at 37°C, the enzymatic reaction was halted by boiling at 100°C for 20 min. The samples were centrifuged at 13,000 rpm for 5 min, and the supernatant was transferred to 800 μl of assay buffer containing 50 mM HEPES-NaOH (pH 7.5), 50 μM Amplex Red, 75 μg/ml D-amino acid oxidase, 54 μg/ml horseradish peroxidase, and 10 mM MgCl_2_. After incubation at 37°C for 60 min, the amount of resorufin was measured at 563 nm using UV-Vis spectrophotometer UVmini-1240 (Shimadzu, Japan).

### Detection of the physical interaction of PG carboxypeptidases with PBPs

Physical interactions between PG carboxypeptidases and PBPs (PBP1a, PBP1b, PBP2, and PBP3) were detected through pull-down experiments. To analyze the pull-down assay, we constructed pBAD-based plasmids expressing C-terminal His-tagged PBPs or non-tagged PBPs. These plasmids were transformed into the MG1655 chromosomal DacC-Flag, DacC(ΔC)-Flag, DacA-Flag, and DacA(ΔC)-Flag strains. The strains were cultured in LB medium at 37°C to an OD_600_ of approximately 0.5. After addition of 1% arabinose, the cells were cultured at 30°C for 3 h. Harvested cells were suspended in 3 ml of binding buffer without 1% sodium n-dodecyl-β-D-maltoside and were disputed twice by a French press at 10,000 psi. After centrifugation at 9,000 × *g* for 10 min at 4°C, the supernatant was mixed with 0.5 ml of the Talon metal affinity resin. After washing the resin three times, the bound proteins were eluted using elution buffer. Total proteins were separated on 4–20% Tris–glycine polyacrylamide gels (KOMA Biotech) and were transferred onto PVDF membranes. The protein levels of DacC, DacC(ΔC), DacA, and DacA(ΔC) were determined using a monoclonal antibody against Flag-tag (Santa Cruz Biotechnology), according to the standard procedure.

To analyze the effect of alkaline pH on the physical interactions between PG carboxypeptidases and PBP1b, ER2566 strains harboring the pET-based plasmids expressing His-tagged DacC, His-tagged DacA, or non-tagged PBP1b were cultured in 100 ml of LB medium at 37°C. When the OD_600_ was approximately 0.5, 1 mM IPTG was added to medium, and then the cells were cultured at 16°C overnight. Harvested cells were suspended in 4.5 ml of binding buffer at pH 8.0 or 9.0, as described above, and were disrupted twice by a French press at 10,000 psi. After centrifugation at 9,000 × *g* for 30 min at 4°C, only the supernatant was mixed with 100 μl of the Talon metal affinity resin in a 1.5-ml tube. After gentle rotation at 4°C and centrifugation at 7,000 rpm for 30 s, the supernatant was deliberately removed. Proteins were bound to the resin by repeated mixing, rotation, centrifugation, and supernatant removal. The resin was washed more than five times with 1 ml binding buffer at pH 8.0 or 9.0. The bound proteins were eluted with elution buffer containing 200 mM imidazole and were separated on 4–20% gradient Tris–glycine polyacrylamide gels (KOMA Biotech). The separated proteins were visualized by staining with Coomassie brilliant blue R.

### Evaluation of cell shape using microscopy

The cells were cultured at 37°C in LB medium at various pH with or without 0.2% arabinose or antibiotics. At an OD_600_ of approximately 0.4, the cells were stained with FM4-64 [N-(3-triethylammoniumpropyl)-4-(*p*-diethylaminophenylhexatrienyl)-pyridinium dibromide] and spotted on a 1% agarose pad prepared in phosphate-buffered saline. The cells were visualized under an Eclipse Ni microscope (Nikon, Japan).

## Funding

This work was supported by research grants from Basic Science Research Program through the National Research Foundation of Korea funded by the Ministry of Education (NRF-2020R1I1A2058026, 2021R1A6A3A01086677, and 2021R1A6A3A01086629) and Korea Institute of Planning and Evaluation for Technology in Food, Agriculture and Forestry (IPET) through High Value-added Food Technology Development Program, funded by Ministry of Agriculture, Food and Rural Affairs (MAFRA) (grant number 322026-3).

## Supplementary materials

### 1. Supplementary Figures

**Figure 3–figure supplement 1.**
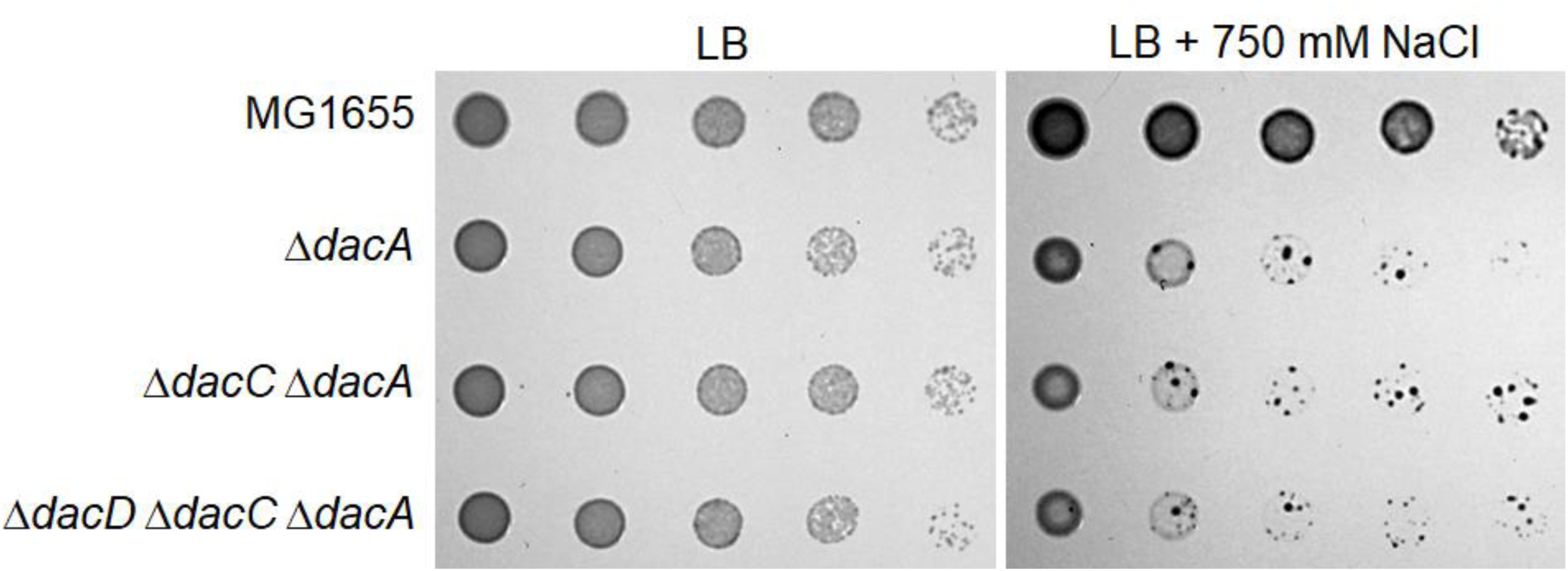
Only DacA is required for overcoming salt stress. The wild-type and indicated mutant cells were serially diluted from 10^8^ to 10^4^ cells/ml in 10-fold steps and spotted onto an LB plate or an LB plate containing 750 mM NaCl.

**Figure 8–figure supplement 1.**
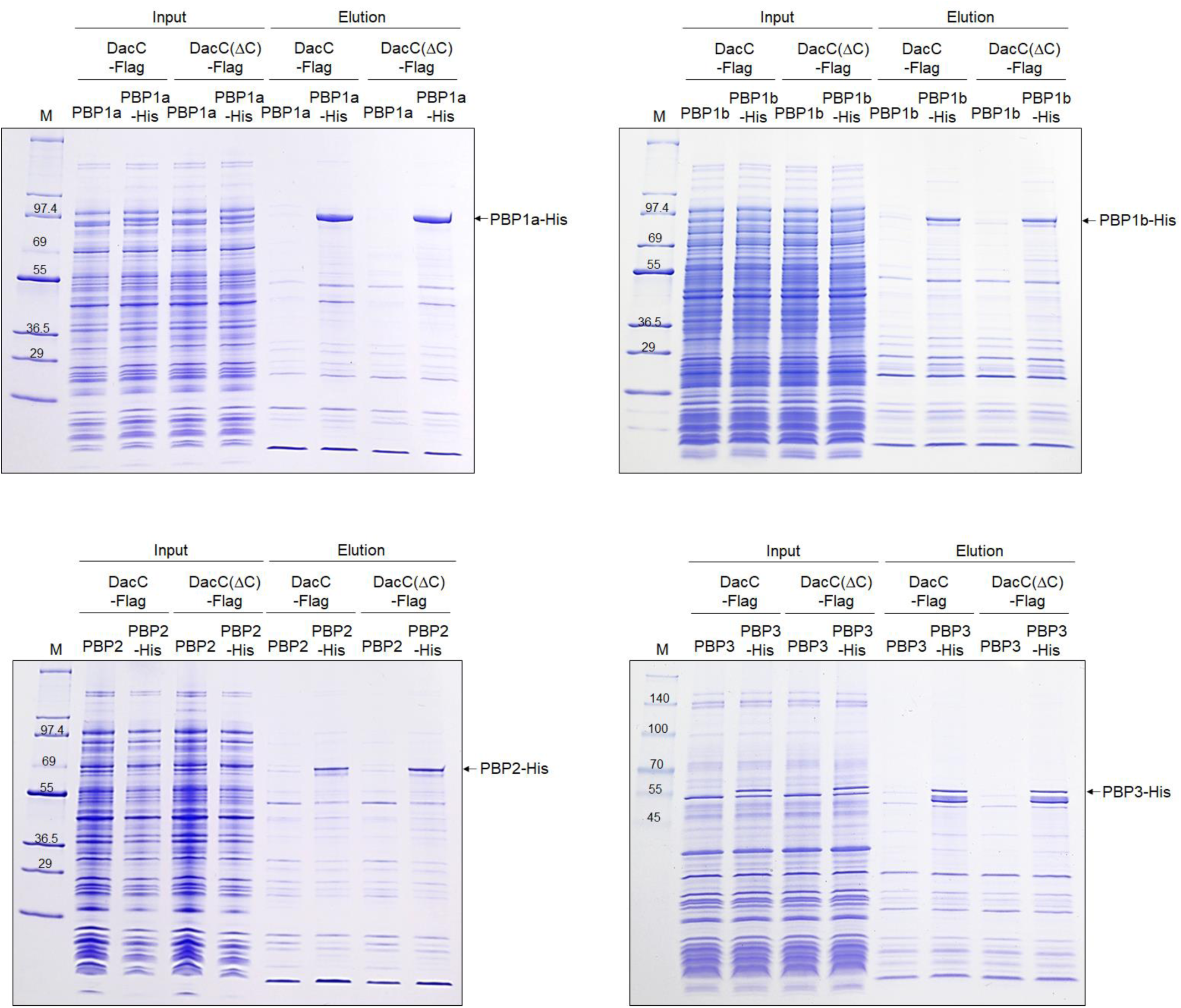
SDS-PAGE gels of pull-down experiments of DacC. The supernatant of MG1655 DacC-Flag or DacC(ΔC)-Flag cells harboring the pBAD-based plasmids expressing C-terminal His-tagged PBPs or non-tagged PBPs was loaded onto Talon metal affinity resin. After pull-down experiments, the amount of input (Input) and output (Elution) total proteins was measured by SDS-PAGE using 4–20% gradient Tris-glycine polyacrylamide gels (KOMA biotech, Korea). Arrows indicate purified His-tagged PBPs. Lane M indicates EzWay Protein Blue MW Marker (KOMA Biotech., Korea) or EzWay Protein PreBlue Ladder (KOMA Biotech., Korea).

**Figure 8–figure supplement 2.**
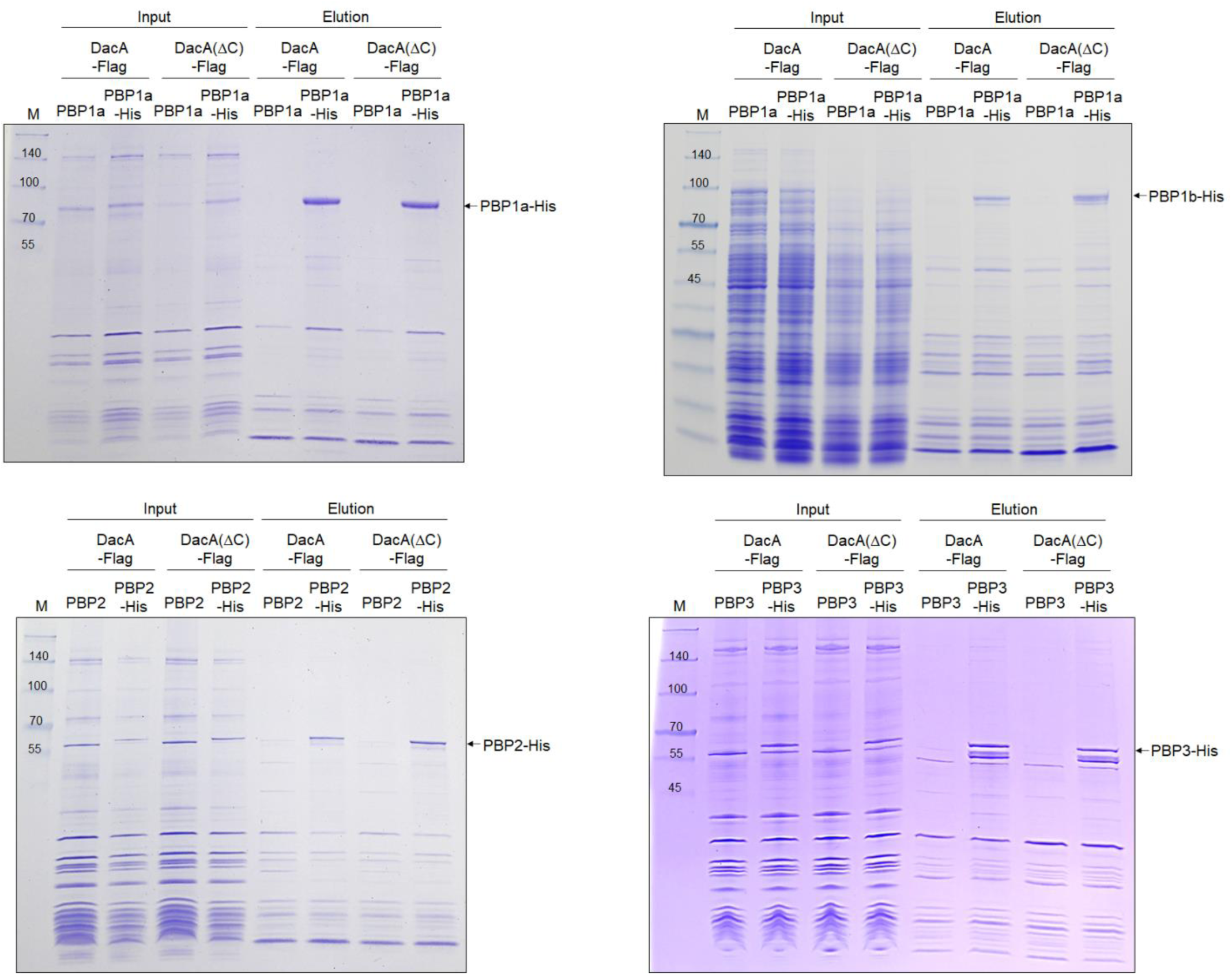
SDS-PAGE gels of pull-down experiments of DacA. The supernatant of MG1655 DacA-Flag or DacA(ΔC)-Flag cells harboring the pBAD-based plasmids expressing C-terminal His-tagged PBPs or non-tagged PBPs was loaded onto Talon metal affinity resin. After pull-down experiments, the amount of input (Input) and output (Elution) total proteins was measured by SDS-PAGE using 4–20% gradient Tris-glycine polyacrylamide gels (KOMA biotech, Korea). Arrows indicate purified His-tagged PBPs. Lane M indicates EzWay Protein PreBlue Ladder (KOMA Biotech., Korea).

**Figure 8–figure supplement 3.**
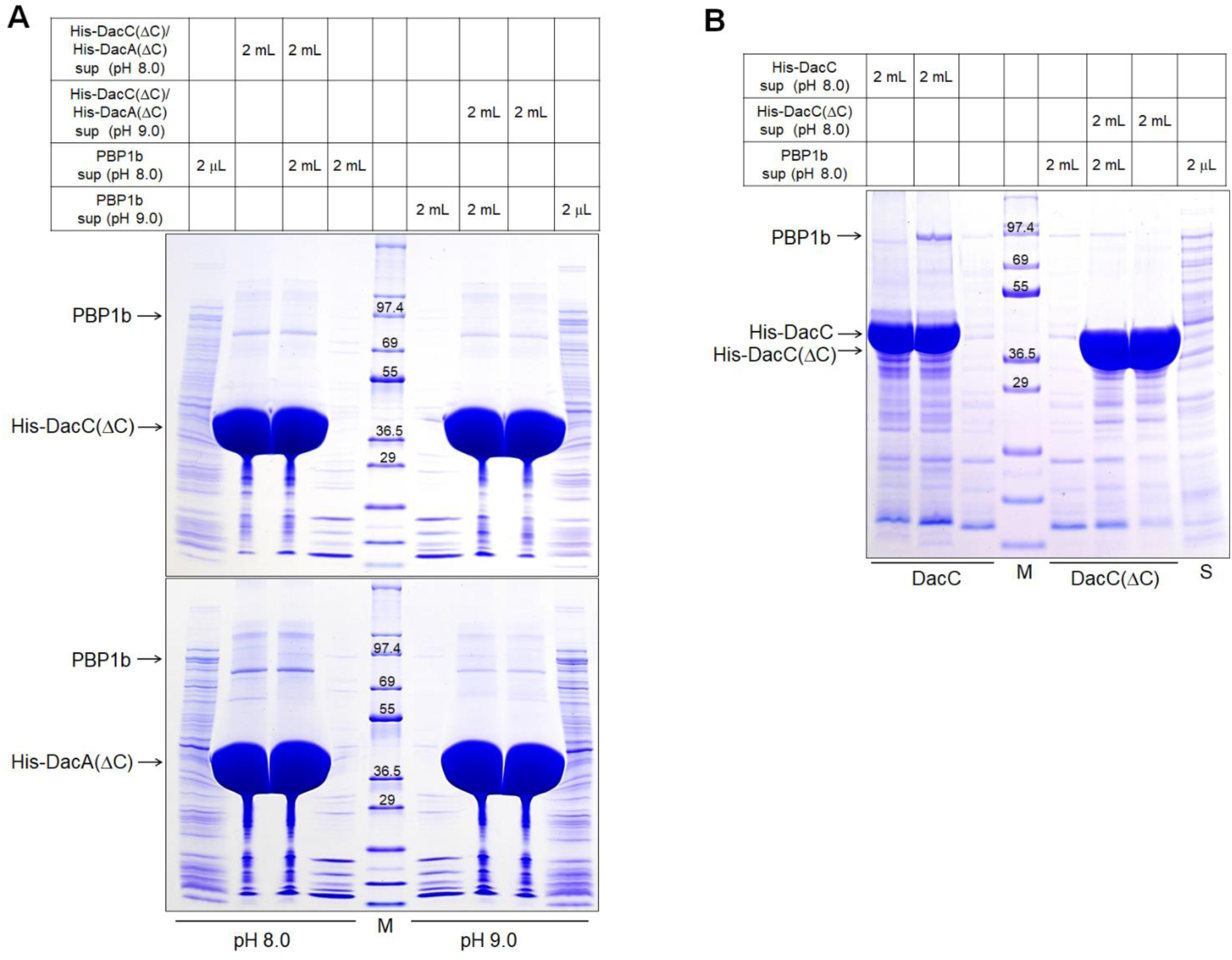
The importance of the C-terminal domain of DacC and DacA for its interaction with PBP1b. (A) PBP1b does not interact with DacC(ΔC) and DacA(ΔC) regardless of pH conditions. The supernatant of ER2566 cells harboring the pET24a plasmid expressing PBP1b was mixed with the supernatant of ER2566 cells harboring pET28a plasmid expressing His-tagged DacC(ΔC) or His-tagged DacA(ΔC) at pH 8.0 or 9.0. After pull-down experiments, eluted proteins were separated on 4–20% gradient Tris-glycine polyacrylamide gels and were visualized by staining with Coomassie brilliant blue R. Lane M indicates EzWay Protein Blue MW Marker (KOMA Biotech., Korea). (B) PBP1b interacts with DacC, but not DacC(ΔC). The supernatant of ER2566 cells harboring the pET24a plasmid expressing PBP1b was mixed with the supernatant of ER2566 cells harboring experiments, eluted proteins were separated on 4–20% gradient Tris-glycine polyacrylamide gels and were visualized by staining with Coomassie brilliant blue R. Lane M indicates EzWay Protein Blue MW Marker (KOMA Biotech., Korea). Lane S indicates the supernatant of ER2566 cells overexpressing PBP1b.

### 2. Supplementary Tables

**Supplementary Table S1.**
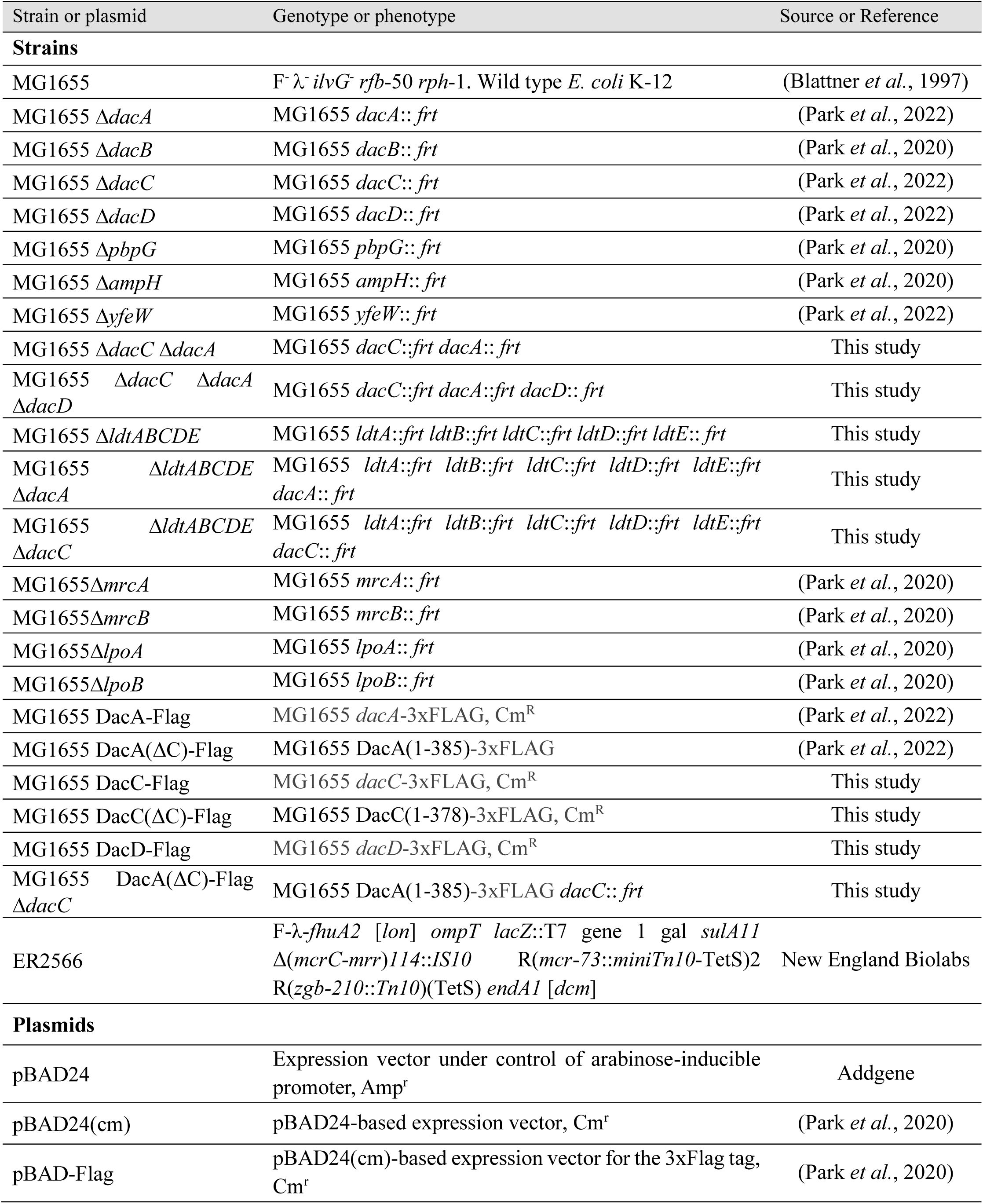
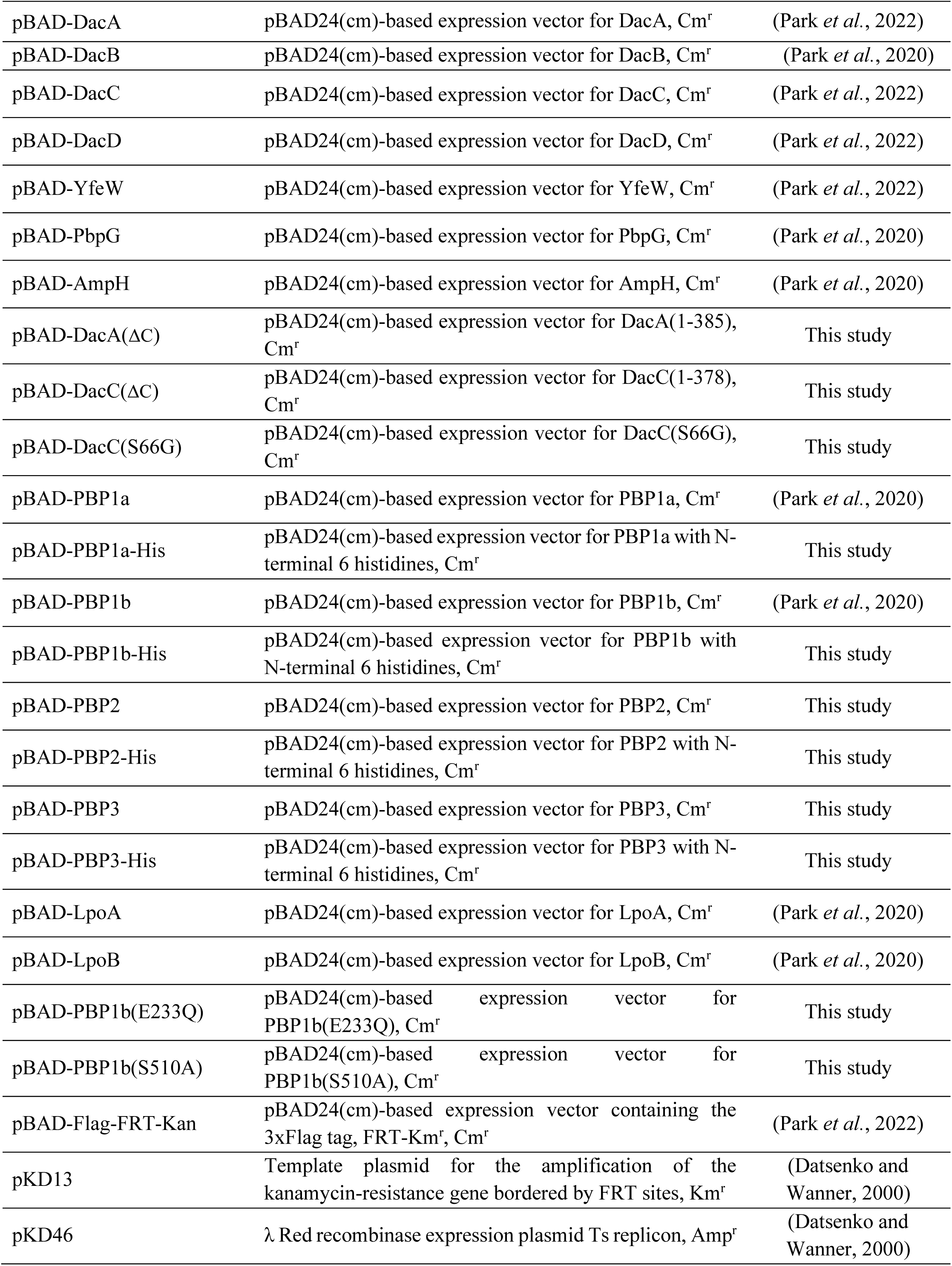
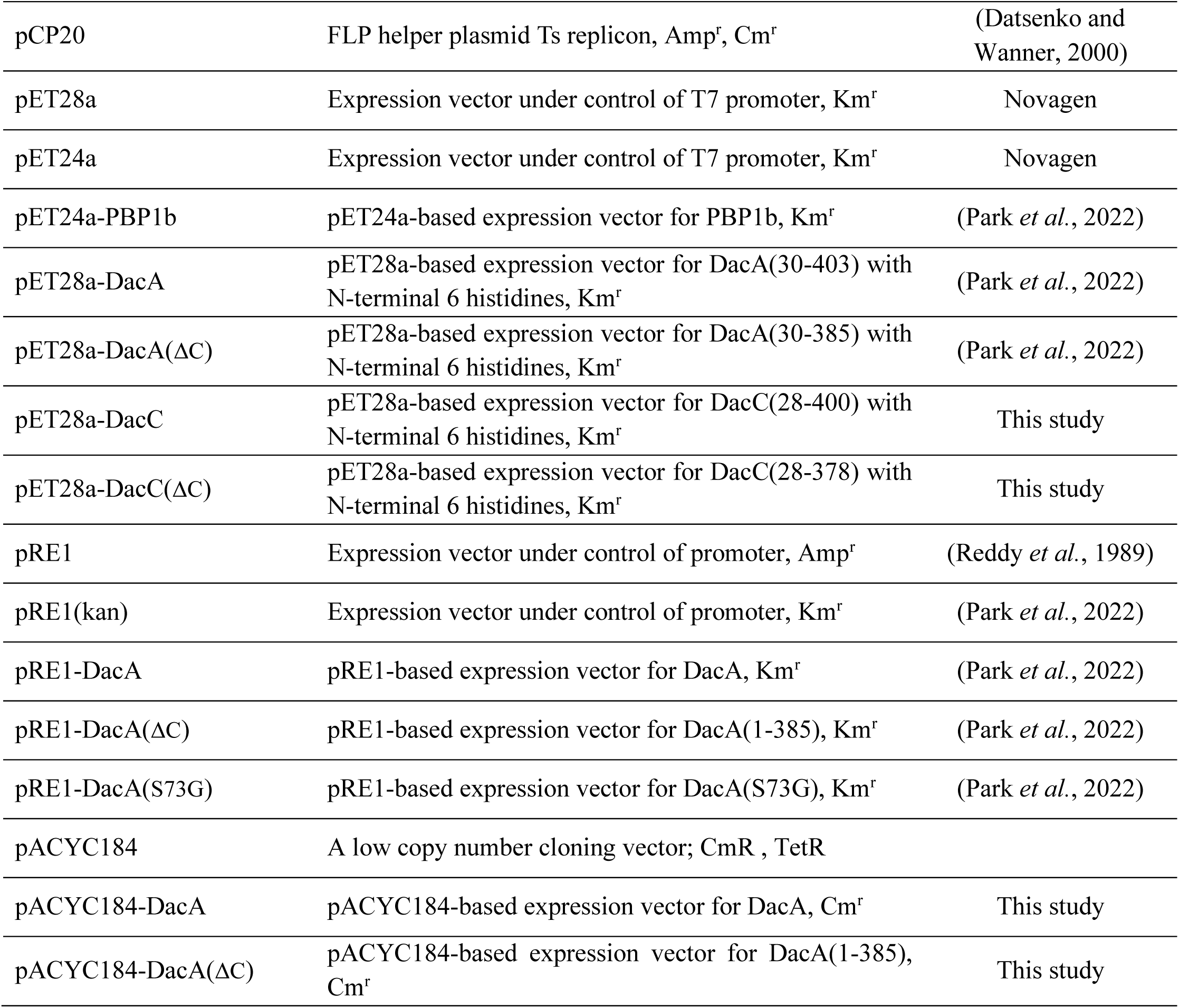
*Escherichia coli* strains and plasmids used in this study.

**Supplementary Table S2.**
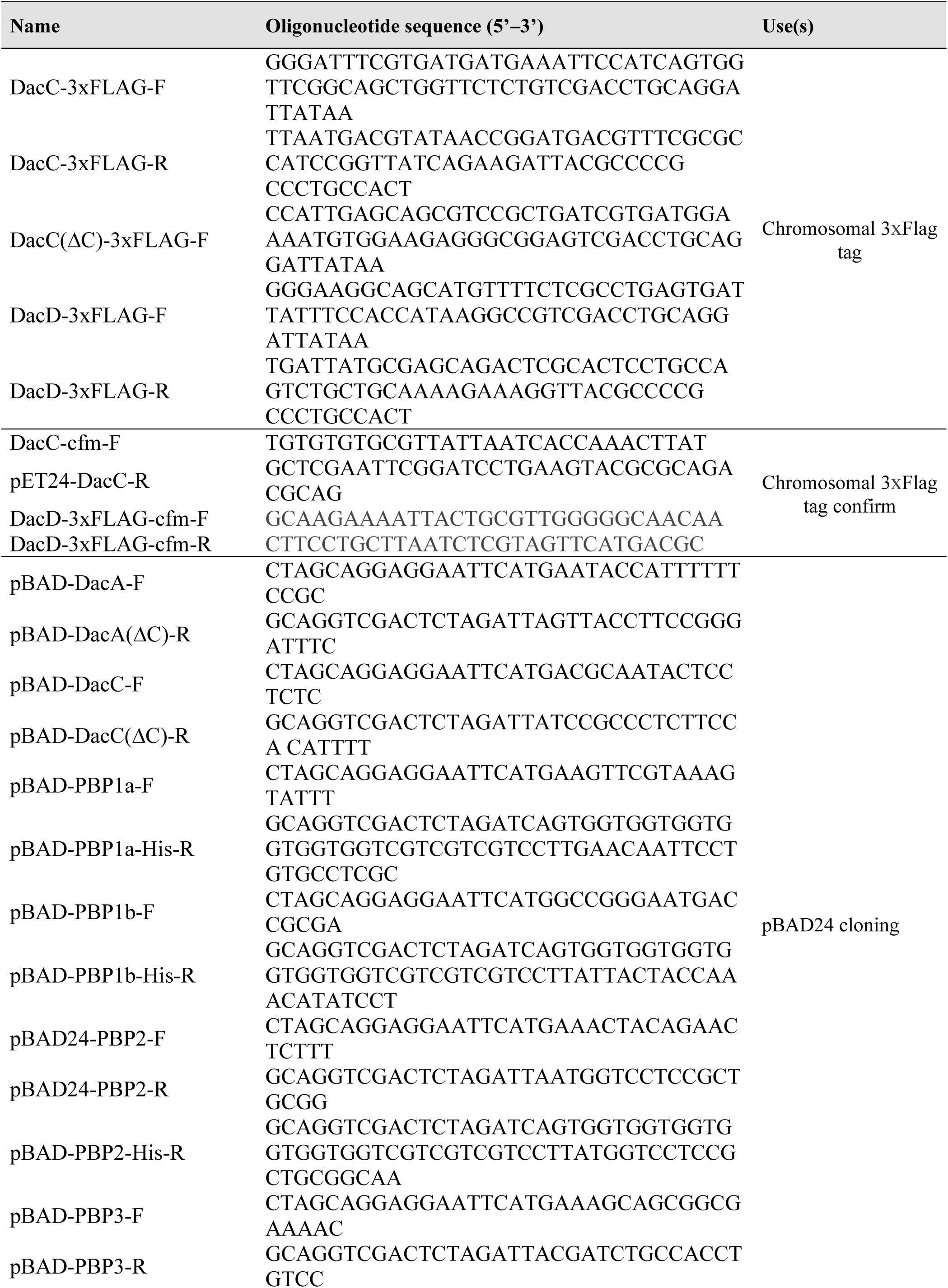
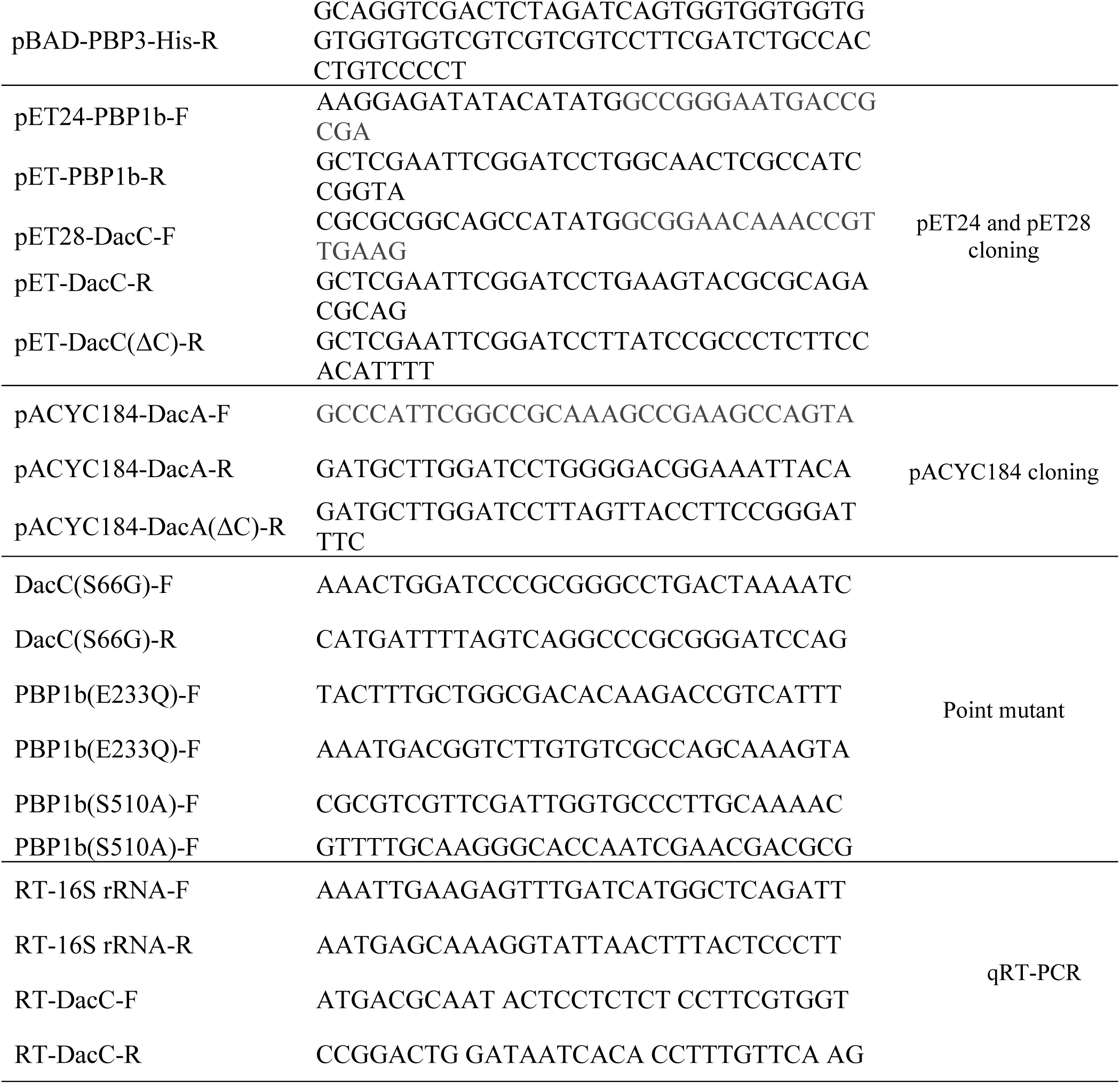
Oligonucleotides used in this study.

## Notes

### Competing Interest Statement

The authors have declared no competing interest.

